# Phase partitioning rules Rab domain formation, growth and identity

**DOI:** 10.1101/2023.04.17.537227

**Authors:** Ana Joaquina Jimenez, Séverine Divoux, Bruno Goud, Franck Perez

## Abstract

Diverse cellular processes are regulated by the formation of specific membrane domains displaying specific lipid and protein compositions. Liquid-liquid phase separation (LLPS) recently emerged as one possible mechanism for their biogenesis, but the examples remain scarce and the impact of LLPS properties on this process is not well established. Rab GTPases are present on all intracellular membranes and play a crucial role in membrane identity, trafficking and compartmentalization. Domain formation is thought to play a central role in Rab functions. Here we show that several Rab partners with common structural characteristics present LLPS properties, some being able to co-condensate or on the contrary presenting immiscible properties. These properties limit the recruitment of Rabs to the membrane sub-domains defined by the condensates of their specific partners. We further show that those LLPS properties control the recruitment of Rab5 to Rabaptin5 condensates ensuring Rab5 functions in regulating endocytic vesicle tethering and fusion. We propose a universal mechanism for Rab domain formation, delimitation, growth and docking based on LLPS properties of Rab partners.

## Introduction

Membrane domains are highly organized regions hosting an intricate network of proteins and lipids and facilitating various critical functions by acting as dynamic platforms for cellular signaling, that regulate communication and coordination between cellular compartments and with the extracellular medium. The importance of membrane domains and their finely-tuned organization extends to critical processes like endocytosis, exocytosis, and the formation and maturation of cellular organelles. Elucidating the mechanisms behind the establishment of these domains is therefore vital for understanding numerous cellular processes and advancing our knowledge of disease mechanisms.

Intracellular membranes involved in traffic consist of a continuum of evolving membranes, which transform into each other. Rab proteins play an essential role in establishing and maintaining the identity state of those membranes . They constitute a large subfamily of small GTPases which are involved in diverse processes, including endocytosis, exocytosis, and organelle trafficking. Rabs serve as master regulators of membrane trafficking, playing a pivotal role in the dynamic movement of vesicles and organelle maturation within the cell. They achieve this by orchestrating the formation of distinct membrane domains that act as molecular signposts that regulate several membrane dynamic processes like binding to molecular motors and ensuring sorting, release and targeting of vesicles, or like mediating vesicle tethering and fusion and therefore promoting organelle growth [5–7].

Most small GTPases undergo post-translational modifications with lipid chains (prenylation) allowing them to anchor to membranes [8,9], and they cycle from soluble to membrane bound depending on their GDP or GTP state. The GDP soluble form high affinity complexes with guanine dissociation inhibitors (GDI) [10,11], which ensure the caging of the lipidic group avoiding membrane anchoring. This cycling is regulated on the one hand by guanine nucleotide exchange factors (GEFs), which ensure the exchange from GDP to GTP and the release from the GDI allowing membrane anchoring. On the other hand, inactivation is mediated by GTPase activating proteins (GAPs), which accelerate the hydrolysis of the GTP into GDP [12,10]. Several examples of Rab GEFs interaction with Rab effectors are known, possibly triggering feed-forward loops of their own recruitment, like for example the Rab5 GEF Rabex5 with the Rab5 effector Rabaptin5 [13,14]. Those GEF/effector interactions have also been suggested to be a general mechanism for domain formation and symmetry breaking [14]. Nevertheless, although feed-forward loops may explain transient local enrichment of Rabs and partners, diffusion is likely to rapidly dilute such clustering, and no mechanism for Rab enrichment containment has been formally proposed.

Among Rabs, the activation of Rab5 by Rabex5/Rabaptin5 complex has been best studied [13,15–17]. Rabaptin5 is a modular protein containing multiple interaction domains: two Rab5 binding domains among which: one specific to Rab5-GTP, one for Rab4 and a another for Rabex5 [15–21]. Despite the existence of two Rab5-binding domains, Rabaptin5 presents higher affinity for Rab5-GTP [13,21] and is most frequently found in complex with Rabex5 [13]. In addition, Rabaptin5 is also found in oligomers with other Rab5 partners like EEA1 [22]. Besides, a possible first step of this domain formation may pass through Rabex5 recruitment to early endosomes (EE) via two distinct Ubiquitin binding domains [23]. Rabaptin5 is essential for Rab5 activation by releasing the autoinhibition of Rabex5 [15,17,18,24]. All these interactions are believed to contribute to a feed-forward loop of Rab5 recruitment, likely promoting biased binding of Rab5-GTP to membranes and promoting domain formation [25,26].

Notably, early endosomes present Rab5 enriched domains [22,27,28] where Rab5 regulates vesicle tethering and fusion [29–33], in a Rabex5-Rabaptin5 complex dependent manner [13,21]. A related role of CapZ in this mechanism through the regulation of actin nucleation has been described [34].

In the past few years, liquid-liquid phase separation (LLPS) has been revealed in biology as a powerful mechanism for the biogenesis and maintenance of membrane-less organelles in cells like nucleoli, P-bodies or stress granules [35], bringing an important contribution to intracellular organization and membrane less compartmentalization. LLPS allows fast molecule storage and release, concentration control and biochemical reaction tuning [36], but also domain formation [37]. Few Rab partners have been described to present phase separation properties, among them the Rab6 effectors ELKS and BICD2. ELKS condensates promote the recruitment of other actors of the polarized membrane associated platforms (PMAPs) near cell edges , and especially the recruitment of Rab6. Properties of LLPS for a partner of Rab8, MICAL-L2 and a partner of Rab3, Rabphilin3A, have also been described [43,44]. *In vitro* studies of Eps15, a potential partner of Rab28 [45], also suggest the implication of Eps15 and Fcho1/2 and their LLPS properties in endocytosis initiation [46]. A study in yeast suggests that Ede1 (the yeast homologue of Eps15) condensates promotes the neoformation and maturation of endocytic sites [47]. Nevertheless, the behavior of the condensates of different Rab effectors have not, to our knowledge, been studied together, as well as their role in defining specific Rab membrane domains.

Here we show that several partners of Rab proteins have phase-separation properties. Those proteins present similar characteristics with long coil-coiled domains and unstructured domains, among them Eps15 and ELKS. Taking the example of Rabaptin5, we show that Rabaptin5 condensates form close to membranes (plasma membrane and endosomal membranes) and that they promote the enrichment of Rab5 and endosome tethering. They also lead to Eps15 condensate fusion, which may enhance the very early steps of endocytosis by promoting the well-established heterotypic fusion between clathrin-coated vesicles and early endosomes [13]. Condensates from partners related to different Rabs present partial or no co-condensation. We propose that phase separation of Rab partners ensure efficient and specific Rab recruitment promoting domain formation and maintenance. These condensates may control also mixing or exclusion between different Rabs depending on the affinity of the involved partners and their capacity or inability to co-condensate.

## Results and discussion

### Rabaptin5 presents LLPS properties sensitive to concentration and time

The multiple interactions of the complex Rabaptin5-Rabex5 with Rab5 is an ideal configuration for network assembly and chain reactions of recruitment [13,21]. The structure of Rabaptin5 predicted by AlphaFold [48] anticipates the presence of long alpha helices and unstructured domains (Fig. 1A) and a disorder analysis done by different algorithms predicts a high disorder score, with a highly disordered large domain in the middle of the protein (Fig. 1B). This, combined with its modular nature , places it as a good candidate to be a scaffold protein for a phase separation phenomenon, which we hypothesized to be a ruling mechanism for Rab5 domain formation.

**Figure 1.**
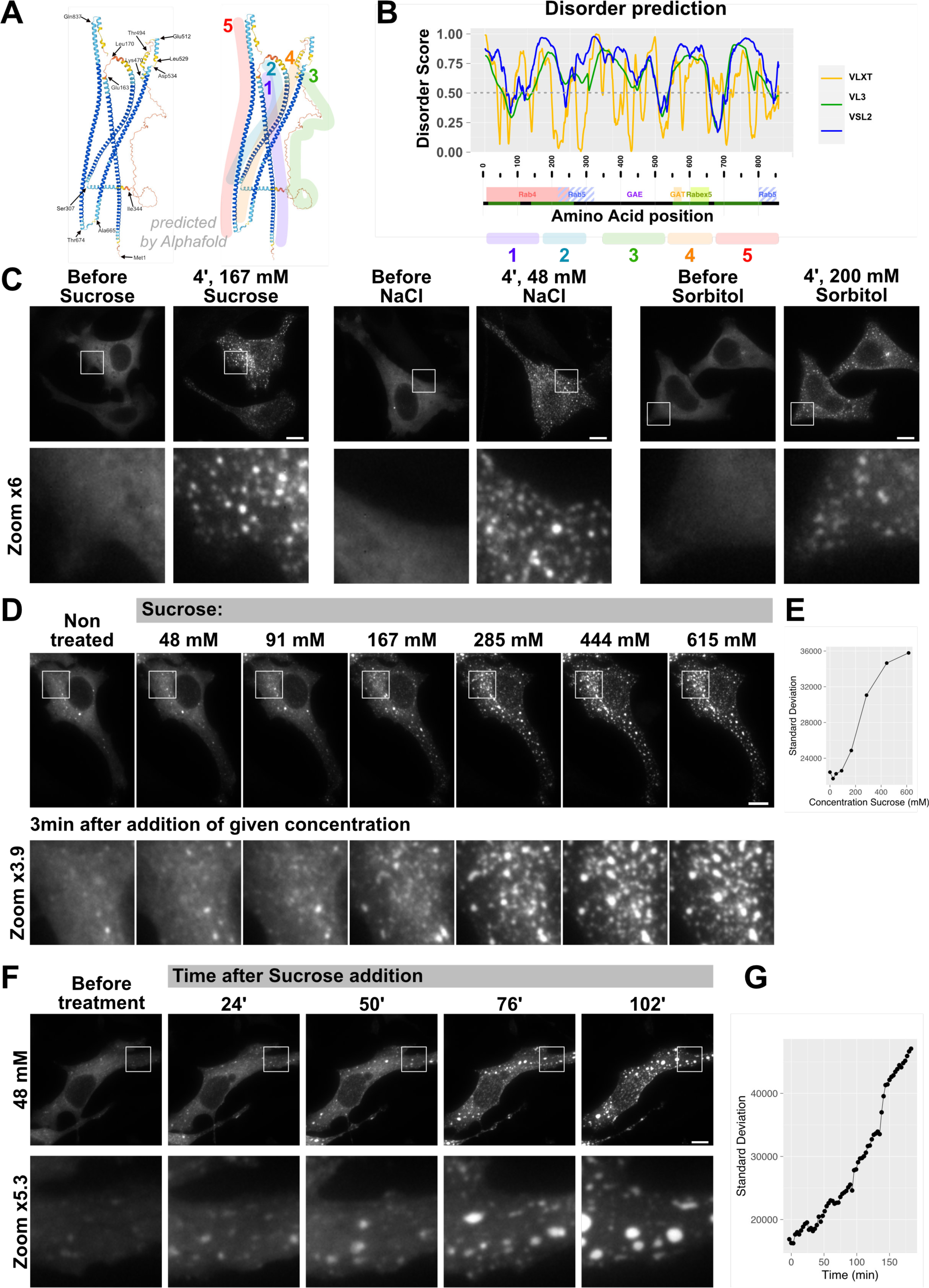
Rabaptin5 present characteristics of phase separating proteins and form puncta upon hyperosmotic treatment in a solute concentration and time dependent manner. **A-** Structure for Rabaptin5 predicted by AlphaFold. **B-** Disorder analysis of Rabaptin5 amino acid sequence using different algorithms: VLXT (yellow), VL3 (green), VSL2 (blue). Dashed line corresponds to a disorder score of 0.5, above which amino acid sequence is considered disordered. **C-** Rabaptin5 forms foci shortly after sucrose, NaCl and Sorbitol treatments in HeLa cells transfected with low levels of EGFP-Rabaptin5. **D-** Foci fluorescent intensity increases with the increase extracellular concentration of sucrose (or NaCl or Sorbitol Fig. 1 SUP2) **E**. as reflected by an increase of the standard deviation of the gray levels plotted against sucrose concentration. **F-** Foci appear at low sucrose concentrations and their intensity is time-dependent. **G-** as reflected by an increase of the standard deviation of the gray levels plotted against time. Scale bars: 10µm.

Thus, we investigated the changes in Rabaptin5 distribution within HeLa cells transfected with low levels of EGFP-Rabaptin5, upon an extemporal increase in concentration using hyperosmotic treatment induced by non-ionic (sucrose and sorbitol) and ionic (NaCl) solutes. We observed that Rabaptin5, initially rather uniformly distributed in the cytoplasm, formed condensates upon an increase in extracellular medium osmolarity (Fig. 1C), even upon a minimal increase of NaCl in the medium (48mM final, 10% increase). EGFP, used as a control soluble protein, did not react to changes in osmolarity nor did CHMP4B known to form puncta upon moderate over-expression (Fig. S1A, B), and shown to oligomerize [49]. Both ionic and non-ionic solutes induced phase separation (although lower concentrations of NaCl were needed, compared to sucrose), suggesting that Rabaptin5 response is due mainly to increased intracellular concentration. This is consistent with previous finding of Rabaptin5 enlarged structures observed upon overexpression, which may correspond to condensates instead of enlarged endosomes proposed by the authors [19].

Because of the higher sensitivity to NaCl addition, we cannot exclude an extra potentiating role of intracellular ion concentration changes on Rabaptin5 phase separation. To limit multi-parameters variations and limit the additional effects that an ionic solute like NaCl may induce besides the changes induced in Rabaptin5 intracellular concentration, we chose to pursue our experiments with sucrose. The size of Rabaptin5 condensates increased with the concentration of sucrose in the extracellular medium as reflected by the increased local intensity variations measured by the standard deviation of the fluorescence within the cell (Fig. 1D, Fig. S1C, E, corresponding graphs in Fig. 1E, Fig. S1D, F). Condensate growth is also time-dependent as low concentrations of sucrose that induced mild or non-detectable LLPS, induced significant condensates after longer incubations (Fig. 1F-G). Of note, this over-time effect of sucrose is surprising given that LLPS laws predict a bi-stable state switch controlled by the existence of a critical concentration of the phase separating molecule. We can envision several explanations for this observation. One could be that the cell is already above the critical concentration of Rabaptin5 and that sucrose is only promoting growth of pre-existent condensates. Another possible explanation is that the cell is able to stabilize long lived condensates that could undergo a gelification-like process as it was described for certain ELKS2 condensates [40].

Importantly, in support to the first hypothesis, many condensates grew from pre-existing Rabaptin5 faint puncta, suggesting that Rabaptin5 small condensates were already present at steady state (Fig. 2A). Such condensates may have been overlooked because in low expressing cells they required adapted acquisition or observation settings (Fig. S1G).

**Figure 2.**
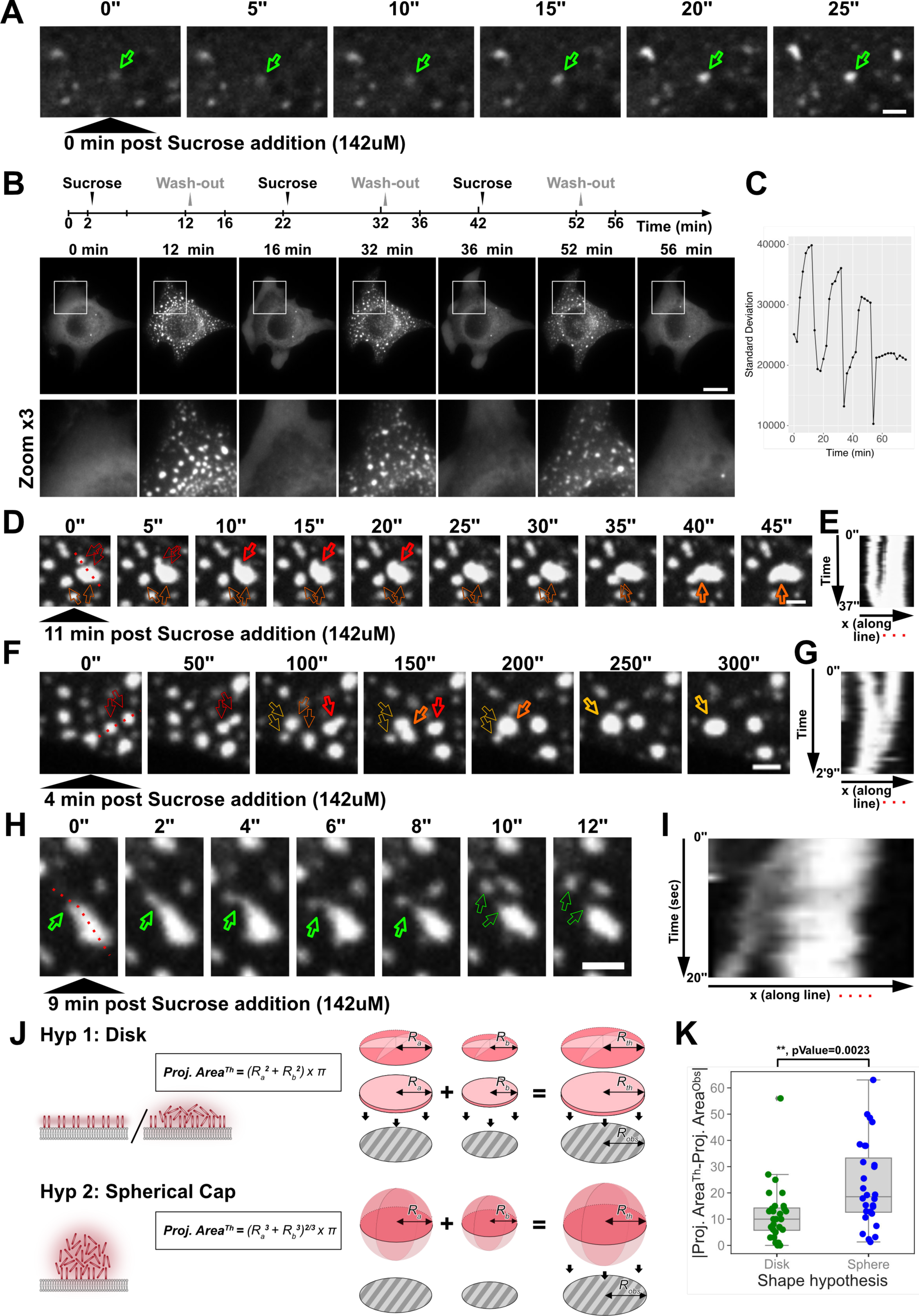
Rabaptin5 condensates behave like seed for larger condensates, are rather flat, reversible and can undergo fusion and fission. **A-** Untreated cells contain Foci capable of growth upon sucrose addition. Empty green arrow points at one of such condensates. **B-** Time-lapse imaging coupled with cycles of addition and removal of sucrose reveal reversibility of condensates as evaluated by the measurement of standard deviation inside cell contour in **C**. **D,F-** Fast time-lapse imaging reveals several events of condensates fusion with different fusion rates. Thin contour empty arrows point at condensates before fusion and thick contour empty arrows point at fused condensate. Different colors point at different fusion events. **E,G-** panels are kymographs corresponding to the dashed lines drew in the left panels on D and F respectively. **H,I-** Time-Lapse imaging and corresponding kymograph reveals occasional fission event. **J,K-** Comparison of observed areas of condensates previous and ulterior to fusion with theoretical areas calculated based on “Disk” and “Sphere” models suggest the condensates have a small height to diameter ratio suggesting “flat” shapes. Scale bars A,D,F,H: 2µm; scale bar B: 10µm.

Because sucrose treatment, as done here, is estimated to increase the concentration of a soluble protein by less than 2-fold, it was surprising that cells with higher concentration of Rabaptin5 did not already present large condensates at the steady state. Indeed, as proposed before, there is a critical concentration (Csat) where condensates should start to form. One likely explanation is that the Csat is already reached in most cells, as suggested by the faint Rabaptin5 patches capable to grow upon sucrose addition. In addition, and as suggested previously, Rabex5 is also needed to create membrane domains [25,26], suggesting that the Csat might need to be calculated for the couple Rabex5-Rabaptin5 and not for Rabaptin5 alone.

### Rabaptin5 condensates presents properties compatible with LLPS

The observed Rabaptin5 condensates presented commonly accepted LLPS characteristics: reversibility (Fig. 2B, C, Movie S1) and capacity to undergo fusion (Fig. 2D-G, Movie S2) and fission (Fig. 2H, I). Rabaptin5 condensates presented frequently irregular shapes, which made us hypothesize that they are not free spherical droplets wandering in the cytoplasm. In order to determine whether Rabaptin5 condensates presented a rather spherical or a rather flat shape, we tested two predictive models for their fusion (Fig. 2J). We imaged cells at high frequency (1 Hz) in order to track condensates over time and follow fusion events. We evaluated the projection area of the fusing condensates before and after fusion and we compared the resulting fusion area of the projections with the theoretical projection areas predicted under the two hypotheses: 1-if the condensates are close to a disk; 2-if the condensates are spheres (Fig. 2J). We compared the absolute value of the difference between the observed area (Proj. Area^Obs^) and the theoretical values (Proj. Area^Th^) given by each model (Fig. 2K). The disk model explained better the observed values of projected condensate areas resulting from fusion events. This suggests that Rabaptin5 condensates are rather flat, close to a 2D condensate or a flatten drop, rather than having a shape close to spherical.

The flat and sometimes irregular shape nature of Rabaptin5 condensates suggest that those condensates could be docked to membranes (see later, Fig. 3C-E, Movie S3). Interestingly, membrane surfaces can potentiate LLPS in several ways and particularly lowering the critical concentration required for LLPS [37,50].

**Figure 3.**
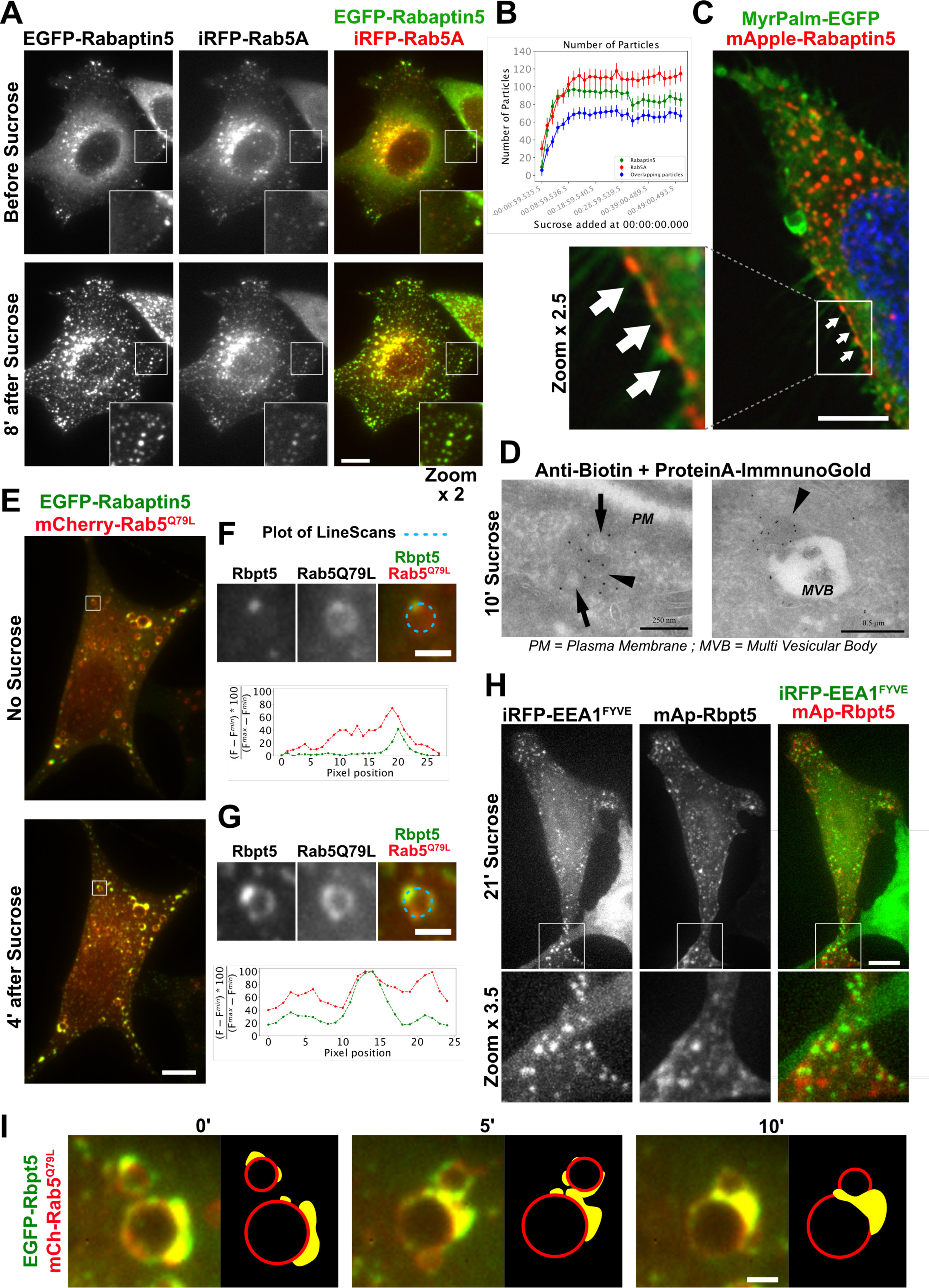
Rabaptin5 form flat condensates docked to membranes which are enriched in Rab5 but not in markers of mature early endosomes. **A-** Z-Stack imaging of cells co-expressing EGFP-Rabaptin5 and mCherry-Rab5 WT condensates reveals enrichment of Rab5 in Rabaptin5 condensates. **B-** Quantification of the number of puncta is quantified after sucrose addition for both proteins, as well as the number of overlapping puncta. **C-** Puncta can be observed at cell edges on cells co-expressing mApple-Rabaptin5 and EGFP-MyrPalm. **D-** Cells were transfected with TurboID-mApple-Rabaptin5 and, 24h hours later, treated with sucrose (147µM) in presence of biotin (40µM) for 10 min, then washed, fixed and processed for cryo-immuno-electron microscopy. Rabaptin5 condensates were detected with anti-biotin antibodies and 10nm Protein-A-gold. Black arrow heads indicate electron-denser condensates labeled with the anti-Biotin. Black arrows point at small vesicles inside the condensates. **E-** Cells transfected with EGFP-Rabaptin5 and Rab5 Q79L mutant present Rabaptin5 condensates docked on enlarged endosomes, capable of growth and Rab5 enrichment upon sucrose treatment as visible on the perimeter scans along enlarged endosome perimeter, indicated by dashed blue lines and plotted in **F** and **G** for before and after sucrose conditions. Intensity profiles were normalized using the minimum between the minimum value found in profile before and after sucrose, and similarly for the maximum, for each channel independently. More examples are available in Figure 3-S1. **H-** Cells expressing mApple-Rabaptin5 and the FYVE domains of EEA1 capable of PI3P binding present juxtaposed but not overlapping puncta. **I-** Enlarged endosomes undergo docking through fusion of Rabaptin5/Rab5Q79L positive condensates. Scale bars A,C,E,H: 10µm; scale bar D-left: 250nm; scale bar D-right: 0.5µm; scale bars F,G,I: 2µm.

### Rabaptin5 condensates promote the recruitment of Rab5 and form preferentially affixed to membranes and at the cell periphery

Rapabtin5 condensates are enriched in Rab5 (Fig. 3A, Fig. S2A, B) and a similar number of Rab5-positive and Rabaptin5-positive condensates appear upon sucrose addition (Fig. 3B). While Rab5 WT (Fig. 3A, Fig. S2A, B) and the GTP-locked mutant Rab5Q79L (Fig. 3E) are efficiently co-condensing, the GTP-binding deficient mutant Rab5 S34N, which inhibits endocytosis and endosome fusion [30–32,51], could not be recruited into Rabaptin5 condensates (Fig. S2C, D) and the GTP binding deficient and prenylation deficient mutant Rab5 N133I [30,31] was only partially recruited (Fig. S2E, F). Like Rab5Q79L, Rab5 N133I likely has a similar conformation than GTP-bound Rab5 and is able to bind directly to Rabaptin5 [30,52]. Taken together, this suggests that Rab5 in its GTP-bound conformation can be recruited to Rabaptin5 condensates but needs to be prenylated to be efficiently enriched.

As observed with other condensates [53,54], Rabaptin5 condensates grow docked to membranes. Fig. 3C and Movie S3 illustrate condensates growing close to the plasma membrane and Fig. 3E close to endosomal membranes (enlarged by the expression of the Rab5 Q79L mutant [32]). We aimed to detect Rabaptin5 condensates using cryo-electron microscopy (cryo-EM) but because of the difficulties to identify morphologically a membrane less element and the rare immunogold labelling usually obtained by classical methods, we decided to amplify the signal by using Rabaptin5 fused to TurboID [55]. This allows a promiscuous biotin ligation to proteins surrounding Rabaptin5 and an enhanced immuno-labelling of Rabaptin5 structures through detection of biotinylated proteins (Fig. S3). Using this approach, we could detect on EM pictures denser and membrane-less structures strongly labelled with anti-biotin, which probably correspond to Rabaptin5 condensates (Fig. 3D). In agreement with fluorescence observations, these dense structures were found affixed to plasma membrane and endosome-like structures. Interestingly, those structures often contained engulfed vesicles.

Finally, Rabaptin5 condensates on enlarged endosomes were enriched in Rab5 and their growth after sucrose addition is accompanied by increased Rab5 signal (Fig. 3E), as shown in the perimeter scans (Fig. 3F, G); other examples of this observation are presented Fig. S4A-D). Interestingly, Rabapatin5 condensates growing on enlarged endosomes was specific to Rab5 enlarged endosomes as Rabaptin5 condensates did not co-localize with enlarged late endosomes induced by the overexpression of the Rab7 Q67L mutant (Fig. S4E).

### Rabaptin5 condensates appear on selective locations/membranes

Rabaptin5 does not decorate the whole perimeter of enlarged Rab5 positive endosomes, (Fig. 3E). In particular, Rabaptin5 condensates are juxtaposed to, but do not co-localize with, clathrin-coated pits or vesicles (Fig. S4F). They also do not co-localize with markers of mature early endosomes such as PI3P (detected by the co-expression of the FYVE domain of EEA1) (Fig. 3H), or Rabenosyn5 (Fig. S4G, H), which is involved in endosomal fusion [56]. This suggests that Rabaptin5 condensates may intervene at the early stages of early endosome formation.

To test this hypothesis, we artificially forced the recruitment Rabaptin5 on Golgi membranes by using Rab6 or a truncated version of Giantin [57] fused to the FRB domain as a trap and the catalytic domain of Rabex5 fused to a FKBP domain as a hook (Fig. S5A, C, B-middle panel, D-middle panel). Upon sucrose addition, we observed that besides condensate formation at the cell periphery, Rabaptin5 condensates also formed in the center of the cells where the Golgi apparatus is located (Fig. S5B-bottom panel, D-bottom panel). This suggests that the presence of Rabaptin5 is sufficient to nucleate condensates.

As shown upon imaging of enlarged endosomes induced by Rab5 Q79L, Rabaptin5 condensates were frequently found in endosome-endosome contact sites, (Fig. 3 I). This suggests that LLPS could be a mechanism for Rab-mediated membrane tethering and therefore promote endosome fusion. This may also explain previous observations of oligomeric complexes containing Rabaptin5 and involved in the tethering of endosomes [22]. The role of Rab5 and partners on endosome fusion has been described before [13,21,29–32] and importantly the Rab5 Q79L mutant, which is not functionally active in terms of GTPase activity, promotes efficiently not only endosome fusion but also stimulates endocytosis [30,32,51]. This shows that Rab5 GTPase activity is not required for those two functions. We propose that LLPS properties of Rabaptin5, and its ability to recruit client proteins like Rab5, are responsible for endocytosis enhancement. This boost would pass through the ability of condensates to tether vesicles together and promote their fusion and growth, in a Rab5 GTPase activity independent manner. Finally, CapZ has been described to recruit Rabaptin5 and other Rab5 effectors on immature early endosomes and promote fusion [34]. Although we could not observe any CapZ enrichment on Rabaptin5 condensates, we do not exclude that condensates could be a mechanism to enrich CapZ and limit actin polymerization on endosomes to promote fusion.

### Other Rab partners co-condensate with Rabaptin5 or form exclusive condensates

Rabaptin5 is highly conserved among the whole animal kingdom but its amino acid sequence is less conserved in insects (*Drosophila melanogaster*) with only ∼25% identity and ∼45% similarity (identity included) (Fig. S6A, C). Despite this divergence, structure prediction reveals striking similarities (Fig. 4A, Fig. S6B). When the amino acid sequence is aligned following domain correspondence based on structure prediction, the score of similarity increases (Fig. S6B, D). When expressed in mammalian cells, the *Drosophila* ortholog of Rabaptin5 (noted Rabep1^Droso^) also displays phase separation upon sucrose addition (Fig. 4B). Condensates grow from pre-existing faint Rabep1^Droso^ -positive patches (Fig. 4C), suggesting that LLPS is a conserved mechanism.

**Figure 4.**
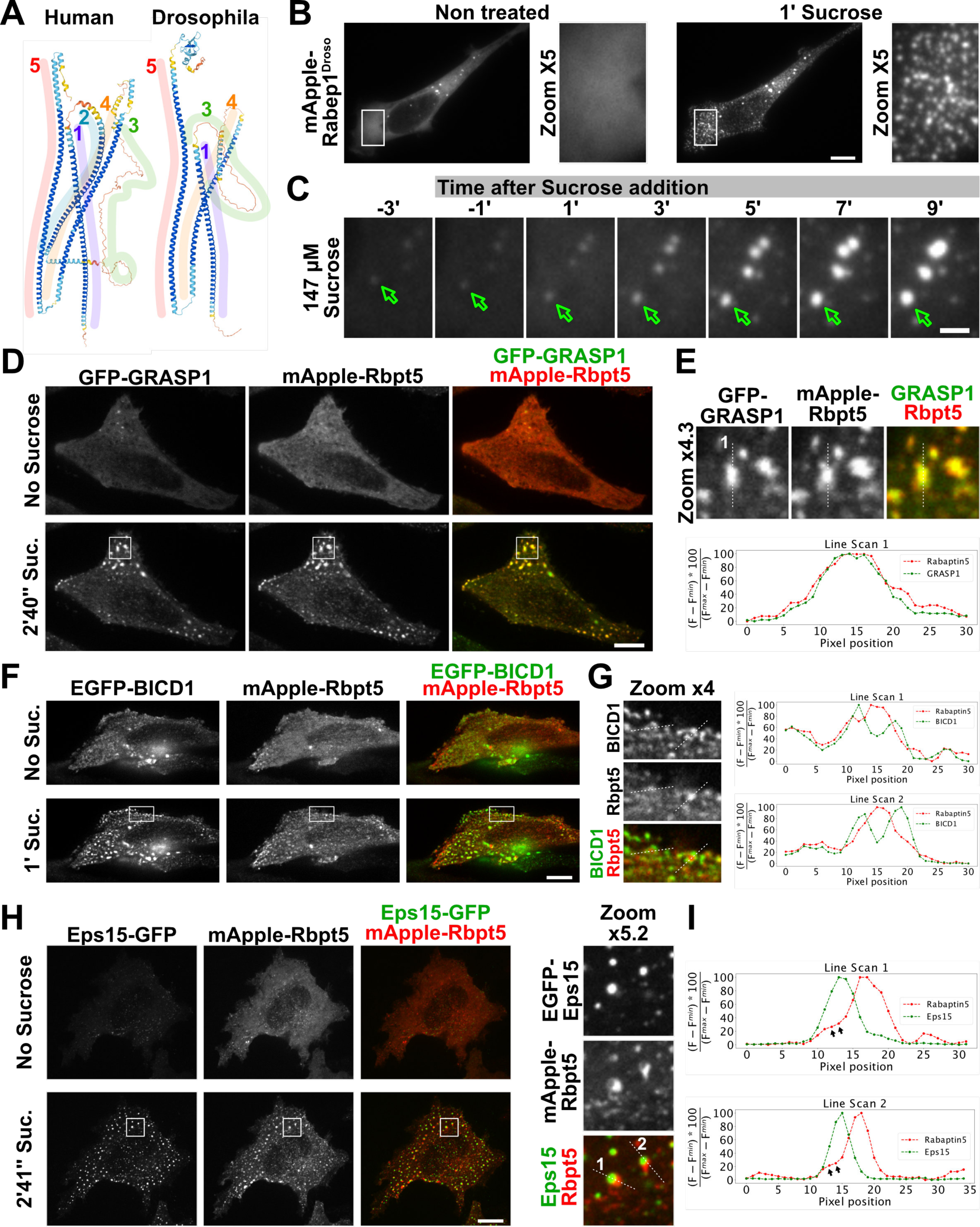
Rabaptin5 LLPS properties are conserved through evolution and are present in many other Rab partners. **A-** Despite high AA sequence divergence (Fig4Sup1) Drosophila and human versions of Rabaptin5 present high predicted structure similarity (Alphafold). **B,C-**Cells expressing the Drosophila version of Rabaptin5 (mApple-Rabep1^Droso^) present growth of condensates from pre-existing structures. **D-** Cells co-expressing EGFP-BICD1 and mApple-Rabaptin5 present exclusive localization as observable by the interspersed line scans pics in **E**. **F,G-** Inversely EGFP-GRASP1 and mApple-Rabaptin5 present overlapped condensates. **H-** Eps15 present an original behaviour with Rabaptin5 condensates hosting Eps15 condensates. **I-** Inflexion pointed by double black arrows in Rabaptin5 line-scan hosts Eps15 intensity pic. Scale bars B,D,F,H: 10µm; scale bar C: 2µm.

Because LLPS properties have been reported before for ELKS and Eps15 [38–42,47], we hypothesize that LLPS could be a mechanism for domain formation common to Rabs. We tested this hypothesis by expressing in HeLa cells various Rab-interacting proteins and found that several of these proteins presented LLPS properties. They all contained similar predicted structural features, namely long alpha helixes and portions of unstructured domains. Those proteins include, in addition to ELKS and Eps15, the Rab6 effector protein BICD1 and the Rab4 partners RUFY1 and GRASP1 [58–60] (Fig. S7). Interestingly, when co-expressed with Rabaptin5, the condensates formed upon sucrose addition presented different degrees of overlapping with Rabaptin5 condensates. Partners of Rabs involved In early endocytosis such as GRASP1 (partner of Rab4, [59]) formed condensates presenting important overlap with Rabaptin5 condensates (Fig. 4D, E). Partners of Rabs not known to be involved in early endocytosis, such as the Rab6 effector proteins ELKS and BICD1, co-localised only partially (ELKS, Fig. S7B), or were completely excluded from Rabaptin5 condensates in the cases of BICD1 (Fig. 4F, G). Interestingly, Rabaptin5 condensates were found to partially co-localise with ELKS at focal additions suggesting once again that Rabaptin5 can localise at or very close to the plasma membrane.

### Rabaptin5 condensates next to Eps15 condensates without mixing

In cells co-expressing mApple-Rabaptin5 and EGFP-Eps15 treated with sucrose, Eps15 condensates appeared as if nestled inside the Rabaptin5 condensates (Fig. 4H, I). This can also be observed in the line scans where Rabaptin5 peaks present an inflexion (indicated by the double black arrows) “nestling” Eps15 peaks (Fig. 4I). Because, Eps15 condensates have been described at the plasma membrane [61,62] and that Rabaptin5 condensates are rather flat and docked to membranes, Eps15 condensate is probably docked to the membrane with a Rabaptin5 condensate shell on top.

Importantly, when we co-expressed mApple-Rabaptin5, EGFP-Eps15 and iRFP-Rab5, we observed that Rab5 presented a clear overlap with Rabaptin5 condensates while it was excluded from Eps15 condensates (Fig. 5A, B, C). This indicates that Rabaptin5 and Eps15 condensates recruit a particular set of partners and can efficiently delimit domains with specific protein composition, such as Rab5 in the case of Rabaptin5 condensates. We then tested how miscible these condensates were. For this, we used a recently published system called CATCHFIRE which allows very fast dimerization of two domains (the ^Fire^Mate and the ^Fire^Tag) upon the addition of a small molecule, the fluorogenic molecule called *match* [63]. We fused Eps15 to the ^Fire^Tag and Rabaptin5 to the ^Fire^Mate in order to induce, upon *match* addition, a rapid and reversible dimerization between Eps15 and Rabaptin5, and evaluate if one protein could integrate the condensates of the other, and if they could form, upon sucrose stimulation, condensates containing both proteins (Fig. 5D-G). The Catch-Fire tagging already promoted condensate formation at steady state in low expressing cells. HMBR addition promoted by itself some enrichment of Rabaptin5 in pre-existing Eps15 condensates and vice-versa (Fig. 5F, G-second panel). Upon sucrose addition, this overlap increased in pre-existing condensates. In addition, previously invisible condensates containing both proteins appeared (Fig. 5F, G-third panel). Upon HMBR wash-out, pre-formed and newly formed condensates remained but the overlap between the two proteins disappeared, suggesting that these two proteins have a natural tendency to unmix.

**Figure 5.**
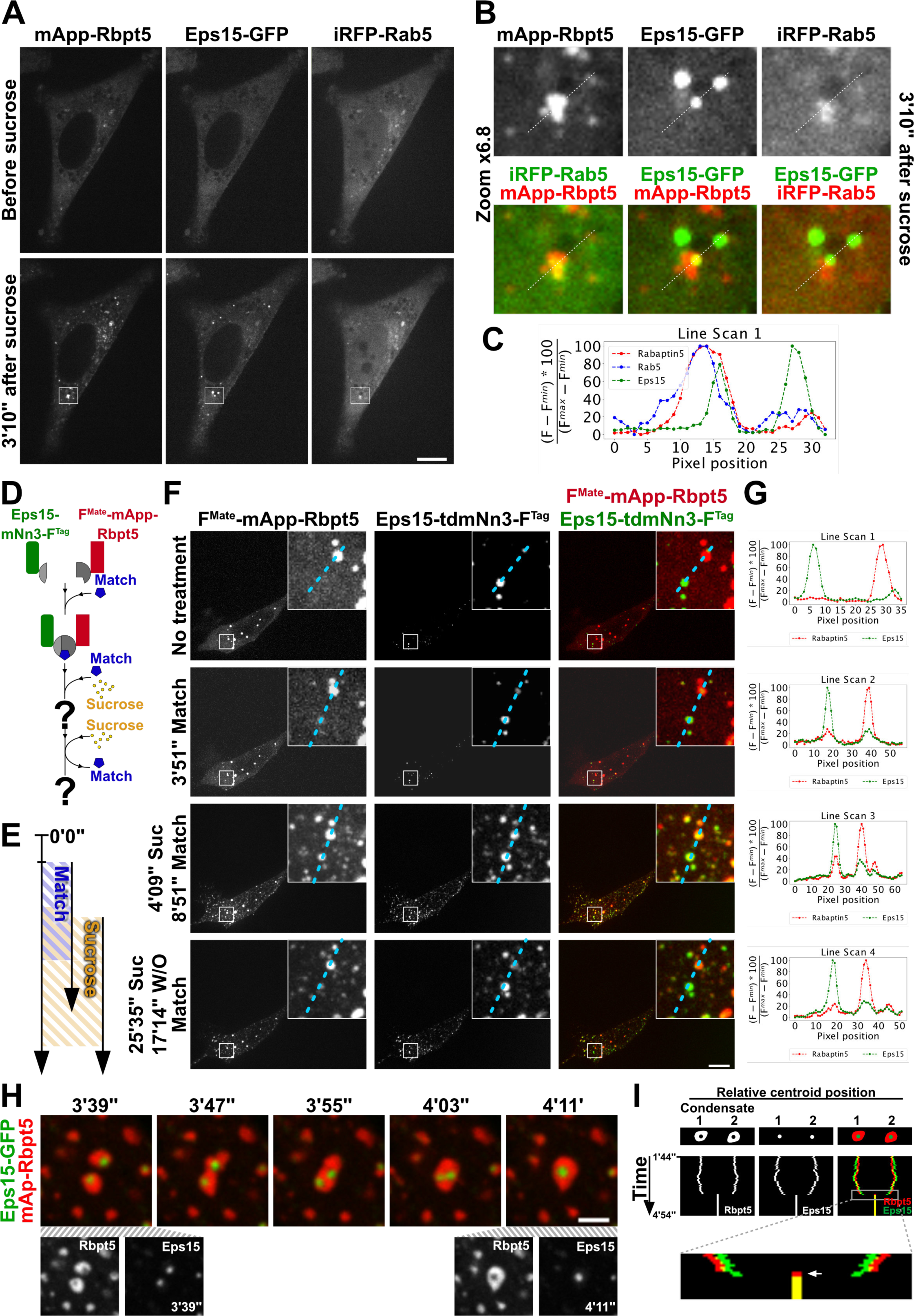
Rab recruitment is condensate specific and Rab effectors present intrinsic properties inducing unmixing. **A,B-** Cells were triple-transfected with mApple-Rabaptin5, EGFP-Eps15 and iRFP-Rab5 show Rab5 co-localization exclusively with Rabaptin5 and not with Eps15. **C-** Line scans corresponding to dashed lines in **B**, show exclusion of Rab5 where Eps15 intensity pics are. **D,E-** Experiment layout of **F** and **G. F,G-** shows some recruitment of Eps15 into Rabaptin5 condensates which disappears shortly after HMBR wash-out. **H-** Fusion of nested Rabaptin5 and Eps15 condensates is homotypic. **I-** Kymograph plots the relative distance between condensates centers of mass over time. Rabaptin5 condensate fusion shortly precedes Eps15 condensate fusion as pointed by white arrow. Scale bars A,F: 10µm; scale bar H: 2µm.

### Rabaptin5 condensates promote Eps15 condensate fusion and growth

Finally, in order to study how partly-imbricated “Eps15-in-Rabaptin5” condensates behave and can influence each other, we co-expressed mApple-Rabaptin5 and EGFP-Eps15.We chose cells with low levels of expression to study the behavior of newly grown condensates upon sucrose addition. We performed fast double color time-lapse imaging (1Hz) in order to be able to track condensates over time. Eps15 and Rabaptin5 condensates moved together but never mixed. Notably, we observed fusion of “Eps15-in-Rabaptin5” partly-imbricated domains (Fig. 5H, Movie S4). Eps15 condensates were able to fuse together inside Rabaptin5 condensates without mixing with Rabaptin5. This process occurs shortly after the fusion of Rabaptin5 condensates, suggesting that the two fusion processes are not independent (Fig. 5I). A tentative hypothesis is that Rabaptin5 condensates promotes Eps15 condensate fusion and growth.

## Conclusion

All together, our results provide strong evidence that Rab proteins interact with proteins with LLPS capacities. Domain formation has been shown to be essential to the function of several Rabs, and in particular for Rab5. We propose that LLPS is the core mechanism that underlie Rab function and drive specific membrane tethering. Additional experiments will be needed to identify each step but our data suggest the following models for domain formation and growth. LLPS properties of Rab effectors and their capacity to co-condensate or to produce immiscible condensates would be a generic mechanism to drive specific and localized recruitment of Rabs, and control domain growth (Fig. 6, panel 1). The involvement of LLPS in Rab domain formation that we propose is also supported by the findings of Bezeljak et al. [26] who showed that the interplay between Rab5 and Rabaptin5-Rabex5 complex behaves like a bistable switch during domain formation. Rab effectors LLPS can thus be used as a bistable switch from a homogeneous state to a phase separation state that would depend on a critical density of protein complexes.

**Figure 6.**
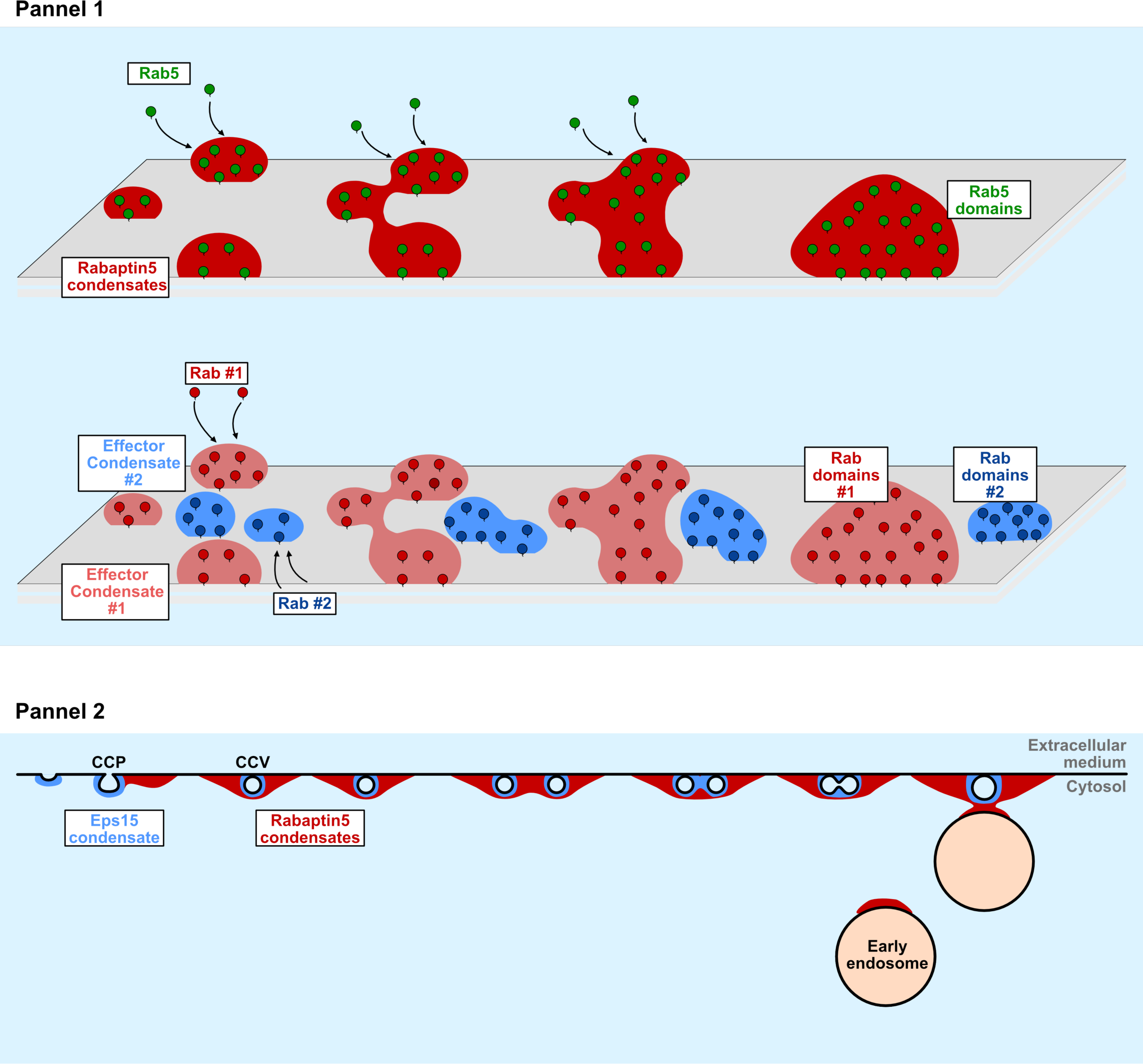
Model. Panel 1-Rab effector condensates mediate specific Rab enrichment, domain delimitation and growth. Panel 2-Rabaptin5 condensates promotes tethering between endocytic vesicles and between endocytic vesicles and early endosomes, ensuring endocytic vesicle maturation.

Interaction between different condensates during growth would then be involved in subsequent membrane maturation steps. Such a maturation has been particularly studied during endocytosis. In particular, we reveal here the existence of nestled bio-condensates that may control the very first steps of clathrin-mediated endocytosis (CME). During those early stages, Eps15 promotes the assembly of clathrin coats that give rise to endocytic vesicles [46,47,61,64,65], which later fuse with early endosomes [13]. Fusion requires Rab5 and Rabaptin5 [66] but its exact mechanism remains unclear. Our results suggest that Rabaptin5 condensates can promote Eps15 condensate fusion while constraining them locally. In parallel, Rabaptin5 condensates can promote tethering between endosomes and we observed that Rabaptin5 condensates contain small vesicles. A tentative hypothesis is that Rabaptin5 condensates mediate homotypic fusion between early endocytic vesicles surrounded by Eps15 condensates, and heterotypic fusion between the late and early endosomes, promoting vesicle growth, and also promoting the maturation of early endosomes (Fig. 6, panel 2).

Finally, Eps15 has been implicated in the recruitment of AP2 and Rab5 has, in turn, been implicated in the AP2 uncoating from clathrin-coated vesicles in partnership with another Rab5 GEF, hRme6 (a.k.a. Rap6, GAPVD1) [67]. It remains to explore whether Rabaptin5 can interact with hRme6 and if Rabaptin5 condensates and their interplay with Eps15 condensates could have a role in this uncoating process.

## Acknowledgements

We thank Marino Zerial, Martin Spiess and Valentina Millarte for sharing Rabaptin5 plasmids. We thank Peter van der Sluijs for Grasp1 and Rufy1 plasmids; Marci Scidmore for BICD1 plasmid from Addgene # 49487; Silvia Corvera for Addgene plasmid # 37538, GFP-Rabenosyn5 ; Ron Vale for Addgene plasmid # 31733. Sergio Grinstein for mCherry-Rab5^Q79L^. Guillaume Montagnac for Clathrin plasmid. Simon de Becco and Mathieu Coppey for iRFP-Rab5^WT^ and iRFP-EEA1^FYVE^ plasmids; Anna Akhmanova for the GFP-ELKS epsilon plasmid; R. Y. Tsien for MyrPalm-EGFP; Justin Taraska for the Eps15-GFP-His from Addgene #170860; Suzanne Pfeffer for the eCFP-Rab7^Q67L^.

We also thank Ahmed ElMarjou and Berengère Ouine for their help with Rabaptin5 purification and Mathieu Coppey for discussion and comments on the manuscript.

We thank the Pict-Ibisa Platform, especially Vincent Fraisier et Chloé Guedj, for microscope trouble shooting.

This work has been supported by the Fondation pour la Recherche Médicale (EQU201903007925), by the Agence Nationale de la Recherche (ANR-19-CE13-0006-03; ANR-20-CE14-0017-02; ANR-19-CE13-0002-03; ANR-23-CE44-0014-02), and has also received support under the program « Investissements d’Avenir » launched by the French Government and implemented by ANR with the references CelTisPhyBio (11-LABX-0038), ANR-10-IDEX-0001-02 PSL. It has also received support from the Labex Cell(n)Scale under the programs “Project Grants” and “Transition Fellowships”.

## Author contributions

A.J.J. designed research, performed most experiments, analyzed the data, drafted the manuscript and provided Labex Cell(n)Scale grants. S.D. performed EM experiments and related preliminary Immunofluorescence tests, provided related figures and commented on the manuscript. B.G. participated to discussions, to reactive founding and edited the manuscript. F.P. funded the project, participated to discussions, followed the advance of the project and edited the manuscript.

## Materials and Methods

### Cell Culture and transfection

HeLa cells were grown in Dulbecco’s modified Eagle medium medium (Gibco BRL) supplemented with 10% fetal bovine serum (Biowest) and 1mM sodium pyruvate, 100 µg/ml penicillin/streptomycin (Invitrogen). Cells were seeded onto 6-well plates, 25mm glass coverslips and transfected in suspension. For expression of the constructs used in this study, HeLa cells were transfected using calcium phosphate precipitates as previously described [68]. For increase in medium osmolarity cells were treated with indicated concentrations with sucrose, sorbitol or NaCl (Merck).

### DNA plasmids

We used the following plasmids received from other teams: eGFP-Rabaptin5 (kind gift from Marino Zerial, MPI-CBG, Dresden, Germany); iRFP-Rab5^WT^ and iRFP-EEA1^FYVE^ (kind gifts from Mathieu Coppey, Institut Curie, Paris, France); mCherry-Rab5^Q79L^ (Addgene #35138, kind gift from Sergio Grinstein, Univ. of Toronto, Toronto, Canada [69]); MyrPalm-EGFP (kind gift from the late Roger Y. Tsien); GFP-GRASP1 and eGFP-Rabip4’/RUFY1 (kind gifts from Peter van der Sluijs, Utrecht University, Utrecht, The Netherlands [58,59]); eGFP-BICD1 (Addgene #49487, kind gift from Marci Scidmore, previously affiliated to Cornell University, Ithaca, U.S.A. [70]); Eps15-GFP-His (Addgene #170860, kind gift from Justin Taraska, NIH, Bethesda, U.S.A. [71]); GFP-Rabenosyn5 (Addgene #37538, kind gift from Sylvia Corvera, UMass Chan Medical School, Worcester, U.S.A. [72]); mCherry-Clathrin Light Chain (kind gift from Guillaume Montagnac, Institut Gustave Roussy, Villejuif, France); GFP-ELKS epsilon (kind gift from Anna Akhmanova, Utrecht University, Utrecht, The Netherlands); eCFP-Rab7^Q67L^ from Canis lupus familiaris, differing from one amino acid from the human ortholog (kind gift from Suzanne Pfeffer, Stanford University School of Medicine, California, U.S.A.). The rest plasmids were constructed as following. eGFP-Rab7^Q67L^: Rab7^Q67L^ was subcloned from eCFP-Rab7^Q67L^ into a pEGFP-C1 plasmid; mApple was a synthesised as a double stranded gene fragment “gBlock” (IDT – Integrated DNA Technologies) and Rabaptin5 was amplified by PCR (using primers 5’-CCAAGCGATCGCTCCGGTGCGCAGCCGGGCCCGGCTTCCCAG-3’ and 5’-GGAGTTAATTAACCATGCGGCCGCTCATGTCTCAGGAAGCTGGTTAATGTCT-3’) and both fragments were cloned into a pA backbone; mCherry-CHMP4B was cloned as previously described [73]; mApple-Rabep1^Droso^ was constructed by extracting Rabep1^Droso^ from a custom synthetic gene (IDT) using humanised codon table, and cloning it into the previously described mApple-Rabaptin5 plasmid instead of the human Rabaptin5; FireMate-mApple-Rabaptin5 was constructed by subcloning the FireMate PCR amplified (using primers 5’-CCTAGCTAGCGCCACCATGGAGCATGTTGCCTTTGGCAGTGAGGA-3’ and 5’-GTGCACCGGTGACCCTCCACCACCTGGAAAGGGCTTTCTTCAAGT-3’) into the previously described mApple-Rabaptin5; the Flag-TurboID-mApple-Rabaptin5 plasmid was built similarly by introducing the Flag-TurboID fragment amplified by PCR (using primers 5’-CCTAGCTAGCATGGATTACAAGGATGACGACGATAAG-3’ and 5’-CGGAGCGATCGCGACCGGTCTTTTCGGCAGACCGCAGAC-3’) into mApple-Rabaptin5; Eps15-tdNano3-FireTag was constructed by cloning a gBlock (IDT) containing a tandem Nano3 couple and a FireTag into the Eps15-GFP-His plasmid replacing the GFP-His part; eGFP-Rab5^N133I^ was built by amplifying eGFP by PCR and using a gBlock for the Rab5^N133I^ mutant and cloning them into a pA backbone; eGFP-Rab5^S34N^ was built similarly; both FRB-Rab6 ires Rab5-FKBP-EGFP and FRB-Mini-Giantin ires-Rabex5-FKBP-EGFP derive from the pIRESneo3 (Clontech-Takara), FRB and FKBP fragments were obtained by digestion, Rabex5 was obtained by PCR (using primers 5’-GAGGCGCGCCATGGGAGGCAGCATTGAAACGGATAGA-3’ and 5’-CATGGAATTCGCCTCAGCTTCTTGCTTCCTGGGAGAGGTCT-3’) and Rab6 and MiniGiantin fragments were purchased as gBlocks (IDT).

### Time lapse fluorescence microscopy

Prior to transfection, HeLa cells were seeded on glass coverslips with a diameter of 25 mm and transfected in suspension. After 20 hours of transfection with the appropriate plasmids, the coverslips were moved to an L-shaped tubing Chamlide from Live Cell Instrument, which was filled with pre-warmed Leibovitz’s medium that is independent of carbonate (manufactured by Invitrogen). At indicated time point, hyperosmotic media was added to the chamber using the tubing. Time-lapse acquisitions were captured at 37 °C within a thermostat-controlled chamber, utilizing an Eclipse 80i microscope from Nikon that is equipped with a spinning disk confocal head from Perkin and a iXon (Andor) camera purchased from Gataca Systems and alternatively a sCMOS Kinetix 22 (Photometrics). Images were acquired using MetaMorph software from Molecular Device, with a 100× objective, with a binning 1 while using the Andor camera and a binning 2 while using the Photometrics camera. Movies destined to condensate tracking were acquired as a single plane and with image frequencies from 1 to 5 Hz.

### Image Analysis and graphics

Custom ImageJ macros were used to mount and analyse the images. Standard deviation was measured at every time point in a region of interest (ROI), adapted to each time point and covering the whole area of the cell. Measurements were performed in the resulting sum of an acquired z-stack covering the entire cell height. Kymographs were performed by tracing by hand a 3-handle spline so that each spline segment crosses through each condensate in the longest possible angle. The spline was then straightened and plotted into a Kymograph. Line scans were performed so that the structure to highlight is at the center of the traced line. Nestled condensate fusion graph was performed as follows: for each time lapse the centroid of each condensate (Rabaptin5 or Eps15) was calculated as well as the relative distance between the two “to fuse” Rabaptin5 condensates or Eps15 condensates. The relative position of each condensate was plotted centred on the vertical axis of the Kymograph. Area projections were measured by choosing “in-focus” condensates and measuring the on-plane area. Calculations were performed using ImageJ. Schemes and figures were done using Affinity Designer Software.

### Catch-Fire assays

To induce dimerization, cell medium was changed for medium containing HMBR. Medium was changed later for medium supplemented with HMBR and sucrose. For wash-out experiments, the cells medium (1ml total), was removed and cells were washed at least 5 times with 1.5ml medium+sucrose, then left in that last mix.

### Cryo-immuno-electron microscopy

HeLa cells were transfected with TurboID-mApple-Rabaptin 5 using calcium phosphate transfection method. After 24h, cells were treated for 10 min with 147 mM sucrose in presence of 40µM Biotin. At the end of the treatment, cells were fixed with 2% paraformaldehyde and 0.2% glutaraldehyde in 0.1M sodium phosphate buffer, pH 7.4. After washing in PBS/Glycin 0,02M, cells were pelleted by centrifugation, embedded in 12% gelatin, cooled in ice and cut into 5-mm^3^ blocks. The blocks were infused over night with 2.3 M sucrose at 4**°**C, frozen in liquid nitrogen and stored until cryo-ultramicrotomy. Sections of 80nm were cut with a diamond knife (Diatome) at **-**120°C using Leica EM-UC7. Ultrathin sections were picked up in a mix of 1.8% methylcellulose and 2.3 M sucrose (1:1) according to Liou *et al*., 1996 and transferred to formvar carbon-coated copper grids.

Immunolabeling was performed as described before (Slot *et al.*, 1991). Cryosections were incubated with a rabbit polyclonal anti-Biotin antibody (Rockland/Tebu-Bio) followed by protein A-10nm gold conjugate (CMC, Utrecht). After labeling, the sections were treated with 1% glutaraldehyde, counterstained with uranyl acetate, and embedded in methyl cellulose uranyl acetate (Slot et al., 1991). The sections were taken up in a wire loop, excess methyl cellulose was sucked into filter paper, and grids were dried before taking them from the loop. Grids were observed on a Tecnai G2 spirit (FEI Company, Eindoven, The Netherlands) equipped with a 4-k CCD camera (Quemesa, Olympus).

### Structure and sequence analysis

Structure predictions were obtained using Aphafold for the following accension numbers (Uniprot): Rabaptin5, Q1527; ELKS, Q8IUD2; BICD1, Q96G01, GRASP1, Q4V328, RUFY1, Q96T51; EPS15, P42566. Rabaptin5 ortholog alignment was performed using Snapgene and the following ascension numbers were evaluated: Homo sapiens isoform1, NP_004694.2; Rattus norvegicus, NP_061997.2; Mus musculus isoform1, NP_062273.2; Bos taurus, NP_001180090.1; Gallus gallus, NP_990388.2; Xenopus tropicalis, NP_001011070.1; Danio renio isoform1, NP_001352427.1; Drosophila melanogaster, NP_611545.1. The domain pairing between human Rabaptin5 and Rabep1^Drosophila^ was handmade based on the Alphafold predicted structure. Disorder domain predictions were made using PONDR (Predictor of Natural Disordered Regions) using VLXT, VL3 and VSL2 algorithms.

### Graphs and statistics

Graphs and statistical tests for dots plots/box plots were performed with Prism 10 (GraphPad) and horizontal bars correspond to mean±SEM. Standard deviation curves were performed using R software. Other time-lapse curves and line scan curves were performed using Jupyter Notebook 6.5.4 (Anaconda Navigator, Python). Curves compiling several time-lapse experiments show at each time-point the mean±SEM. Given the big variability in fluorescence intensity among cells and condensates, curves showing the standard deviation are shown independently for each cell, as well as curves destined to show the coherence between two channels.

## Supplementary information Legends

**Figure S1. LLPS is not a characteristic of all proteins, and the magnitude of the reaction of Rabaptin5 depends on extracellular medium solute concentration and Rabaptin5 intracellular concentration.**

**A-** EGFP does not present Foci formation upon sucrose addition, **B-** neither does CHMP4B which is known to present Foci when overexpressed. **C, E-** Foci fluorescent intensity increases with the increase extracellular concentration of NaCl or Sorbitol respectively, as reflected by an increase of the standard deviation of the gray levels plotted against the concentration of **D-** NaCl or **F-** Sorbitol. Images shown were taken 3 minutes after the addition of medium supplemented with the indicated solute concentration. **G-** The magnitude of foci formation increases with the amount of Rabaptin5 expression but adapted acquisition parameters. All cells were acquired the same day, with the same laser power and exposure so that fluorescence intensity is comparable. We present here cells with up to 15-fold difference in intensity. Because of the high difference in signal intensity, cells are presented with different ranges of gray levels (Look-Up tables or ”LUT”) to be able to observe condensates in all of them. Look-up Tables. Labelled in red allow the observation of foci in very low-expressing cells. Scale bars: 10µm.

**Figure S2. Rab5 mutations impairing prenylation or GTP binding also prevent Rab5 from enriching in Rabaptin5 condensates.**

Time-lapse imaging of cells co-expressing Rabaptin5 and Rab5 and respective particle quantification (**A,B-** WT, **C,D-** prenylation deficient N133I, or **E,F-** GTP binding deficient S34N) reveals decreased Rab5 enrichment on Rabaptin5 condensates as well as a decrease in the number of observed puncta in Rab5 mutants. Scale bars: 10µm.

**Figure S3. Upon treatment with sucrose and biotin, TurboID-mApple-Rabaptin5 is able to efficiently biotinylate its surroundings within condensates.**

**A-** Scheme illustrating the use of Rabaptin5 fused to the promiscuous biotin ligase Turbo-ID, which allows biotinylation of proteins in Rabaptin5 surroundings and allows ImmunoGold signal amplification. **B-** HeLa cells were transfected with TurboID-mApple-Rabaptin 5 and treated with sucrose (147µM) in presence of biotin (40µM) for 10 min, then washed and fixed. Cells were then processed for cryo-immuno-electron microscopy (Fig. 3D) or stained using neutravidin-A488. White arrows point at biotinylated condensates.

**Figure S4. Rabaptin5 condensates localize close to membranes and is accompanied by an enrichment in Rab5 but do not co-localize with markers of mature early endosomes nor with early markers of endocytosis**

**A-D** Cells transfected with EGFP-Rabaptin5 and mCherry-Rab5 Q79L mutant present Rabaptin5 condensates docked on enlarged endosomes, capable of growth and Rab5 enrichment upon sucrose treatment. **A,C-** Close-up on several enlarged endosomes followed before and after sucrose. Dashed blue curves marking the perimeter scans plotted in **B** and **D. B,D-** Straighten perimeter scans of enlarged endosomes before and after sucrose respectively. Like in Figure 3, intensity profiles were normalized using the minimum between the minimum value found in profile before and after sucrose, and similarly for the maximum, for each channel independently. **E-** Cells transfected mApple-Rabaptin5 and EGFP-Rab7 Q67L mutant does not present docking of Rabaptin5 condensates on Rab7 positive enlarged endosomes. **F-** Co-expression of Rabaptin5 and clathrin light-chain reveals Rabaptin5 condensate appearance and some increase in intensity on clathrin puncta, as well as some juxtaposition between the two markers but no co-localization. **G-** Time-lapse imaging of cells co-expressing mApple-Rabaptin5 and GFP-Rabenosyn5 condensates reveals juxtaposition but not co-localization between the two proteins upon sucrose addition. Zoom 1 shows central particles co-labeled with Rabaptin5 and Rabenosyn5. Zoom 2 shows newly showing peripheral particles labeled with Rabaptin5 and little or no Rabenosyn5. **H-** Puncta quantification shows particle number increase for Rabaptin5 but not for Rabenosyn5. It also reveals little overlapping between Rabenosyn5 and Rabaptin5 puncta. Scale bars E-G: 10µm; scale bars A,C: 2µm.

**Figure S5. Artificial indirect recruitment of Rabaptin5 on “foreign” membranes potentiate Rabaptin5 condensate formation.**

**A,C-** Experiment layouts. **B,D-** Rabaptin5 artificially recruited at the Golgi is able to nucleate condensates . Scale bars: 10µm.

**Figure S6. Rabaptin5 is evolutionary conserved in sequence among animals and in structure compared to insects.**

**A-** Table shows the amino acid sequence identities or similarities between Human Rabaptin5 and other species versions. **B-** Alphafold 3D structure prediction of Human and Drosophila versions. **C-** Overall alignment of amino-acid sequences (SnapGene). **D-** Alignment region by region defined by structure prediction in **B**.

**Figure S7. Other Rab partners presenting common 3D features with Rabaptin (long alpha-helixes and long unstructured domains) are also able to undergo LLPS.**

Cells were transfected with indicated Rab partners and treated with medium supplemented with 147mM sucrose. **A,C-F-** Alphafold predicted structures and pictures before and after sucrose are shown, for ELKS, BICD1, RUFY1 (a.k.a. Rabip4’), GRASP1 and Eps15 respectively. **B-** Cells co-expressing mApple-Rabaptin5 and EGFP-ELKS present partial co-localisation. Scale bars: 10µm

**Movie S1. Rabaptin5 condensates forming upon Sucrose addition are reversible.**

HeLa cells were transfected with EGFP-Rabaptin5 24h before the acquisition with a 2min time-lapse under a spinning disk microscope. Sucrose was added at 2, 22 and 42 minutes, and respective wash-outs were performed at 13, 33 and 53 minutes.

**Movie S2. Rabaptin5 condensates are able to undergo fusion.**

HeLa cells were transfected with EGFP-Rabaptin5 24h before the acquisition with a 1 second time-lapse under a spinning disk microscope. Sucrose was added at 18 seconds after the start of the movie.

**Movie S3. Rabaptin5 condensates can appear at the edges of the cell.**

HeLa cells were transfected with EGFP-Rabaptin5 24h before the acquisition with a 1 second time-lapse under a spinning disk microscope. Sucrose was added at 20 seconds after the start of the movie.

**Movie S4. Fusion of nested Rabaptin5 and Eps15 condensates is homotypic.**

HeLa cells were transfected with mApple-Rabaptin5 and Eps15-GFP 24h before the acquisition with a 1 second time-lapse under a spinning disk microscope. Sucrose was added at 28 seconds after the start of the movie. Several events of fusion between nestled condensates are observable.

## Bibliography

1. Barr FA: Rab GTPases and membrane identity: Causal or inconsequential? Journal of Cell Biology 2013, 202:191–199.

2. Mizuno-Yamasaki E, Rivera-Molina F, Novick P: GTPase Networks in Membrane Traffic. Annual Review of Biochemistry 2012, 81:637–659.

3. Pfeffer SR: Rab GTPase regulation of membrane identity. Curr Opin Cell Biol 2013, 25:414–419.

4. Stenmark H: Rab GTPases as coordinators of vesicle traffic. Nat Rev Mol Cell Biol 2009, 10:513–525.

5. Banworth MJ, Li G: Consequences of Rab GTPase dysfunction in genetic or acquired human diseases. Small GTPases 2018, 9:158–181.

6. Homma Y, Hiragi S, Fukuda M: Rab family of small GTPases: an updated view on their regulation and functions. FEBS J 2021, 288:36–55.

7. Miserey-Lenkei S, Bousquet H, Pylypenko O, Bardin S, Dimitrov A, Bressanelli G, Bonifay R, Fraisier V, Guillou C, Bougeret C, et al.: Coupling fission and exit of RAB6 vesicles at Golgi hotspots through kinesin-myosin interactions. Nat Commun 2017, 8:1254.

8. Wang M, Casey PJ: Protein prenylation: unique fats make their mark on biology. Nat Rev Mol Cell Biol 2016, 17:110–122.

9. Ghomashchi F, Zhang X, Liu L, Gelb MH: Binding of prenylated and polybasic peptides to membranes: affinities and intervesicle exchange. Biochemistry 1995, 34:11910– 11918.

10. Cherfils J, Zeghouf M: Regulation of small GTPases by GEFs, GAPs, and GDIs. Physiol Rev 2013, 93:269–309.

11. Sasaki T, Kikuchi A, Araki S, Hata Y, Isomura M, Kuroda S, Takai Y: Purification and characterization from bovine brain cytosol of a protein that inhibits the dissociation of GDP from and the subsequent binding of GTP to smg p25A, a ras p21-like GTP-binding protein. J Biol Chem 1990, 265:2333–2337.

12. Bos JL, Rehmann H, Wittinghofer A: GEFs and GAPs: critical elements in the control of small G proteins. Cell 2007, 129:865–877.

13. Horiuchi H, Lippé R, McBride HM, Rubino M, Woodman P, Stenmark H, Rybin V, Wilm M, Ashman K, Mann M, et al.: A novel Rab5 GDP/GTP exchange factor complexed to Rabaptin-5 links nucleotide exchange to effector recruitment and function. Cell 1997, 90:1149–1159.

14. Goryachev AB, Leda M: Autoactivation of small GTPases by the GEF-effector positive feedback modules. F1000Res 2019, 8:F1000 Faculty Rev-1676.

15. Delprato A, Lambright DG: Structural basis for Rab GTPase activation by VPS9 domain exchange factors. Nat Struct Mol Biol 2007, 14:406–412.

16. Lauer J, Segeletz S, Cezanne A, Guaitoli G, Raimondi F, Gentzel M, Alva V, Habeck M, Kalaidzidis Y, Ueffing M, et al.: Auto-regulation of Rab5 GEF activity in Rabex5 by allosteric structural changes, catalytic core dynamics and ubiquitin binding. Elife 2019, 8.

17. Zhang Z, Zhang T, Wang S, Gong Z, Tang C, Chen J, Ding J: Molecular mechanism for Rabex-5 GEF activation by Rabaptin-5. Elife 2014, 3.

18. Delprato A, Merithew E, Lambright DG: Structure, exchange determinants, and family-wide rab specificity of the tandem helical bundle and Vps9 domains of Rabex-5. Cell 2004, 118:607–617.

19. Kälin S, Hirschmann DT, Buser DP, Spiess M: Rabaptin5 is recruited to endosomes by Rab4 and Rabex5 to regulate endosome maturation. Journal of Cell Science 2015, 128:4126–4137.

20. Lippé R, Horiuchi H, Runge A, Zerial M: Expression, purification, and characterization of Rab5 effector complex, rabaptin-5/rabex-5. Methods Enzymol 2001, 329:132–145.

21. Stenmark H, Vitale G, Ullrich O, Zerial M: Rabaptin-5 is a direct effector of the small GTPase Rab5 in endocytic membrane fusion. Cell 1995, 83:423–432.

22. McBride HM, Rybin V, Murphy C, Giner A, Teasdale R, Zerial M: Oligomeric complexes link Rab5 effectors with NSF and drive membrane fusion via interactions between EEA1 and syntaxin 13. Cell 1999, 98:377–386.

23. Penengo L, Mapelli M, Murachelli AG, Confalonieri S, Magri L, Musacchio A, Di Fiore PP, Polo S, Schneider TR: Crystal structure of the ubiquitin binding domains of rabex-5 reveals two modes of interaction with ubiquitin. Cell 2006, 124:1183–1195.

24. Parkinson G, Roboti P, Zhang L, Taylor S, Woodman P: His domain protein tyrosine phosphatase and Rabaptin-5 couple endo-lysosomal sorting of EGFR with endosomal maturation. Journal of Cell Science 2021, 134:jcs259192.

25. Cezanne A, Lauer J, Solomatina A, Sbalzarini IF, Zerial M: A non-linear system patterns Rab5 GTPase on the membrane. eLife 2020, 9.

26. Bezeljak U, Loya H, Kaczmarek B, Saunders TE, Loose M: Stochastic activation and bistability in a Rab GTPase regulatory network. Proc Natl Acad Sci U S A 2020, 117:6540–6549.

27. Franke C, Repnik U, Segeletz S, Brouilly N, Kalaidzidis Y, Verbavatz J, Zerial M: Correlative single-molecule localization microscopy and electron tomography reveals endosome nanoscale domains. Traffic 2019, 20:601–617.

28. Sönnichsen B, De Renzis S, Nielsen E, Rietdorf J, Zerial M: Distinct Membrane Domains on Endosomes in the Recycling Pathway Visualized by Multicolor Imaging of Rab4, Rab5, and Rab11. Journal of Cell Biology 2000, 149:901–914.

29. Gorvel J-P, Chavrier P, Zerial M, Gruenberg J: rab5 controls early endosome fusion in vitro. Cell 1991, 64:915–925.

30. Hoffenberg S, Sanford JC, Liu S, Daniel DS, Tuvin M, Knoll BJ, Wessling-Resnick M, Dickey BF: Biochemical and functional characterization of a recombinant GTPase, Rab5, and two of its mutants. J Biol Chem 1995, 270:5048–5056.

31. Li G, Barbieri MA, Colombo MI, Stahl PD: Structural features of the GTP-binding defective Rab5 mutants required for their inhibitory activity on endocytosis. J Biol Chem 1994, 269:14631–14635.

32. Stenmark H, Parton RG, Steele-Mortimer O, Lütcke A, Gruenberg J, Zerial M: Inhibition of rab5 GTPase activity stimulates membrane fusion in endocytosis. EMBO J 1994, 13:1287–1296.

33. Wang Y, Roche O, Yan MS, Finak G, Evans AJ, Metcalf JL, Hast BE, Hanna SC, Wondergem B, Furge KA, et al.: Regulation of endocytosis via the oxygen-sensing pathway. Nat Med 2009, 15:319–324.

34. Wang D, Ye Z, Wei W, Yu J, Huang L, Zhang H, Yue J: Capping protein regulates endosomal trafficking by controlling F-actin density around endocytic vesicles and recruiting RAB5 effectors. eLife 2021, 10:e65910.

35. Banani SF, Lee HO, Hyman AA, Rosen MK: Biomolecular condensates: organizers of cellular biochemistry. Nat Rev Mol Cell Biol 2017, 18:285–298.

36. Lyon AS, Peeples WB, Rosen MK: A framework for understanding the functions of biomolecular condensates across scales. Nat Rev Mol Cell Biol 2021, 22:215–235.

37. Snead WT, Gladfelter AS: The Control Centers of Biomolecular Phase Separation: How Membrane Surfaces, PTMs, and Active Processes Regulate Condensation. Mol Cell 2019, 76:295–305.

38. Sala K, Corbetta A, Minici C, Tonoli D, Murray DH, Cammarota E, Ribolla L, Ramella M, Fesce R, Mazza D, et al.: The ERC1 scaffold protein implicated in cell motility drives the assembly of a liquid phase. Sci Rep 2019, 9:13530.

39. Jin G, Lin L, Li K, Li J, Yu C, Wei Z: Structural basis of ELKS/Rab6B interaction and its role in vesicle capturing enhanced by liquid-liquid phase separation. Journal of Biological Chemistry 2023, 0.

40. Liang M, Jin G, Xie X, Zhang W, Li K, Niu F, Yu C, Wei Z: Oligomerized liprin-α promotes phase separation of ELKS for compartmentalization of presynaptic active zone proteins. Cell Rep 2021, 34:108901.

41. McDonald NA, Fetter RD, Shen K: Assembly of synaptic active zones requires phase separation of scaffold molecules. Nature 2020, 588:454–458.

42. Wu X, Ganzella M, Zhou J, Zhu S, Jahn R, Zhang M: Vesicle Tethering on the Surface of Phase-Separated Active Zone Condensates. Molecular Cell 2021, 81:13–24.e7.

43. Sakane A, Yano T-A, Uchihashi T, Horikawa K, Hara Y, Imoto I, Kurisu S, Yamada H, Takei K, Sasaki T: JRAB/MICAL-L2 undergoes liquid-liquid phase separation to form tubular recycling endosomes. Commun Biol 2021, 4:551.

44. Yang L, Wei M, Wang Y, Zhang J, Liu S, Liu M, Wang S, Li K, Dong Z, Zhang C: Rabphilin-3A undergoes phase separation to regulate GluN2A mobility and surface clustering. Nat Commun 2023, 14:379.

45. Ying G, Boldt K, Ueffing M, Gerstner CD, Frederick JM, Baehr W: The small GTPase RAB28 is required for phagocytosis of cone outer segments by the murine retinal pigmented epithelium. J Biol Chem 2018, 293:17546–17558.

46. Day KJ, Kago G, Wang L, Richter JB, Hayden CC, Lafer EM, Stachowiak JC: Liquid-like protein interactions catalyse assembly of endocytic vesicles. Nat Cell Biol 2021, 23:366– 376.

47. Kozak M, Kaksonen M: Condensation of Ede1 promotes the initiation of endocytosis. eLife 2022, 11:e72865.

48. Jumper J, Evans R, Pritzel A, Green T, Figurnov M, Ronneberger O, Tunyasuvunakool K, Bates R, Žídek A, Potapenko A, et al.: Highly accurate protein structure prediction with AlphaFold. Nature 2021, 596:583–589.

49. Teis D, Saksena S, Emr SD: Ordered assembly of the ESCRT-III complex on endosomes is required to sequester cargo during MVB formation. Dev Cell 2008, 15:578–589.

50. Zhao YG, Zhang H: Phase Separation in Membrane Biology: The Interplay between Membrane-Bound Organelles and Membraneless Condensates. Developmental Cell 2020, 55:30–44.

51. Li G, Stahl PD: Structure-function relationship of the small GTPase rab5. Journal of Biological Chemistry 1993, 268:24475–24480.

52. Strick DJ, Francescutti DM, Zhao Y, Elferink LA: Mammalian suppressor of Sec4 modulates the inhibitory effect of Rab15 during early endocytosis. J Biol Chem 2002, 277:32722–32729.

53. Case LB, Zhang X, Ditlev JA, Rosen MK: Stoichiometry controls activity of phase separated clusters of actin signaling proteins. Science 2019, 363:1093–1097.

54. Huang WYC, Alvarez S, Kondo Y, Lee YK, Chung JK, Lam HYM, Biswas KH, Kuriyan J, Groves JT: A molecular assembly phase transition and kinetic proofreading modulate Ras activation by SOS. Science 2019, 363:1098–1103.

55. Branon TC, Bosch JA, Sanchez AD, Udeshi ND, Svinkina T, Carr SA, Feldman JL, Perrimon N, Ting AY: Efficient proximity labeling in living cells and organisms with TurboID. Nat Biotechnol 2018, 36:880–887.

56. Nielsen E, Christoforidis S, Uttenweiler-Joseph S, Miaczynska M, Dewitte F, Wilm M, Hoflack B, Zerial M: Rabenosyn-5, a Novel Rab5 Effector, Is Complexed with Hvps45 and Recruited to Endosomes through a Fyve Finger Domain. J Cell Biol 2000, 151:601– 612.

57. van Unen J, Reinhard NR, Yin T, Wu YI, Postma M, Gadella TWJ, Goedhart J: Plasma membrane restricted RhoGEF activity is sufficient for RhoA-mediated actin polymerization. Sci Rep 2015, 5:14693.

58. Fouraux MA, Deneka M, Ivan V, van der Heijden A, Raymackers J, van Suylekom D, van Venrooij WJ, van der Sluijs P, Pruijn GJM: Rabip4’ is an effector of rab5 and rab4 and regulates transport through early endosomes. Mol Biol Cell 2004, 15:611–624.

59. Hoogenraad CC, Popa I, Futai K, Martinez-Sanchez E, Wulf PS, van Vlijmen T, Dortland BR, Oorschot V, Govers R, Monti M, et al.: Neuron specific Rab4 effector GRASP-1 coordinates membrane specialization and maturation of recycling endosomes. PLoS Biol 2010, 8:e1000283.

60. Ivan V, Martinez-Sanchez E, Sima LE, Oorschot V, Klumperman J, Petrescu SM, Sluijs P van der: AP-3 and Rabip4’ Coordinately Regulate Spatial Distribution of Lysosomes. PLOS ONE 2012, 7:e48142.

61. Mayers JR, Wang L, Pramanik J, Johnson A, Sarkeshik A, Wang Y, Saengsawang W, Yates JR, Audhya A: Regulation of ubiquitin-dependent cargo sorting by multiple endocytic adaptors at the plasma membrane. Proc Natl Acad Sci U S A 2013, 110:11857–11862.

62. Torrisi MR, Lotti LV, Belleudi F, Gradini R, Salcini AE, Confalonieri S, Pelicci PG, Di Fiore PP: Eps15 is recruited to the plasma membrane upon epidermal growth factor receptor activation and localizes to components of the endocytic pathway during receptor internalization. Mol Biol Cell 1999, 10:417–434.

63. Bottone S, Joliot O, Cakil ZV, El Hajji L, Rakotoarison L-M, Boncompain G, Perez F, Gautier A: A fluorogenic chemically induced dimerization technology for controlling, imaging and sensing protein proximity. Nat Methods 2023, 20:1553–1562.

64. Ma L, Umasankar PK, Wrobel AG, Lymar A, McCoy AJ, Holkar SS, Jha A, Pradhan-Sundd T, Watkins SC, Owen DJ, et al.: Transient Fcho1/2⋅Eps15/R⋅AP-2 Nanoclusters Prime the AP-2 Clathrin Adaptor for Cargo Binding. Dev Cell 2016, 37:428–443.

65. Sengar AS, Wang W, Bishay J, Cohen S, Egan SE: The EH and SH3 domain Ese proteins regulate endocytosis by linking to dynamin and Eps15. EMBO J 1999, 18:1159–1171.

66. Rubino M, Miaczynska M, Lippé R, Zerial M: Selective membrane recruitment of EEA1 suggests a role in directional transport of clathrin-coated vesicles to early endosomes. J Biol Chem 2000, 275:3745–3748.

67. Semerdjieva S, Shortt B, Maxwell E, Singh S, Fonarev P, Hansen J, Schiavo G, Grant BD, Smythe E: Coordinated regulation of AP2 uncoating from clathrin-coated vesicles by rab5 and hRME-6. J Cell Biol 2008, 183:499–511.

68. Jordan M, Schallhorn A, Wurm FM: Transfecting mammalian cells: optimization of critical parameters affecting calcium-phosphate precipitate formation. Nucleic Acids Res 1996, 24:596–601.

69. Bohdanowicz M, Balkin DM, De Camilli P, Grinstein S: Recruitment of OCRL and Inpp5B to phagosomes by Rab5 and APPL1 depletes phosphoinositides and attenuates Akt signaling. MBoC 2012, 23:176–187.

70. Moorhead AR, Rzomp KA, Scidmore MA: The Rab6 effector Bicaudal D1 associates with Chlamydia trachomatis inclusions in a biovar-specific manner. Infect Immun 2007, 75:781–791.

71. Prasai B, Haber GJ, Strub M-P, Ahn R, Ciemniecki JA, Sochacki KA, Taraska JW: The nanoscale molecular morphology of docked exocytic dense-core vesicles in neuroendocrine cells. Nat Commun 2021, 12:3970.

72. Navaroli DM, Bellvé KD, Standley C, Lifshitz LM, Cardia J, Lambright D, Leonard D, Fogarty KE, Corvera S: Rabenosyn-5 defines the fate of the transferrin receptor following clathrin-mediated endocytosis. Proc Natl Acad Sci U S A 2012, 109:E471–480.

73. Jimenez AJ, Maiuri P, Lafaurie-Janvore J, Divoux S, Piel M, Perez F: ESCRT Machinery Is Required for Plasma Membrane Repair. Science 2014, 343:1247136.

