## Supplementary Material for "Phase partitioning rules Rab domain formation, growth and identity"

### Supplemental data

### Supplemental Figures

### Figure S1

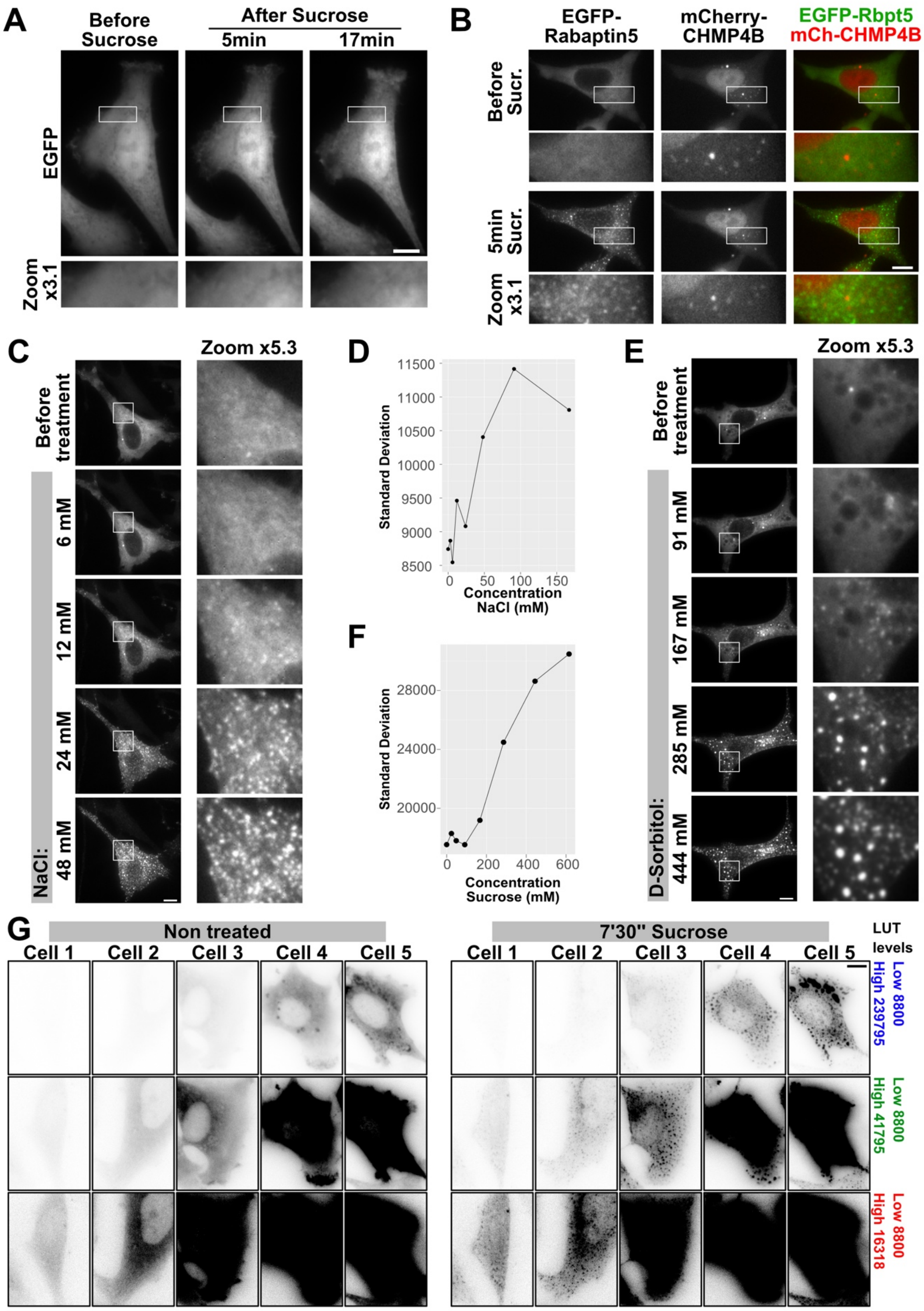


**Figure S1. LLPS is not a characteristic of all proteins, and the magnitude of the reaction of Rabaptin5 depends on extracellular medium solute concentration and Rabaptin5 intracellular concentration.**

**A-** EGFP does not present Foci formation upon sucrose addition, **B-** neither does CHMP4B which is known to present Foci when overexpressed. **C, E-** Foci fluorescent intensity increases with the increase extracellular concentration of NaCl or Sorbitol respectively, as reflected by an increase of the standard deviation of the gray levels plotted against the concentration of **D-** NaCl or **F-** Sorbitol. Images shown were taken 3 minutes after the addition of medium supplemented with the indicated solute concentration. **G-** The magnitude of foci formation increases with the amount of Rabaptin5 expression but adapted acquisition parameters. All cells were acquired the same day, with the same laser power and exposure so that fluorescence intensity is comparable. We present here cells with up to 15-fold difference in intensity. Because of the high difference in signal intensity, cells are presented with different ranges of gray levels (Look-Up tables or ”LUT”) to be able to observe condensates in all of them. Look-up Tables. Labelled in red allow the observation of foci in very low-expressing cells. Scale bars: 10µm.

**Figure S2**

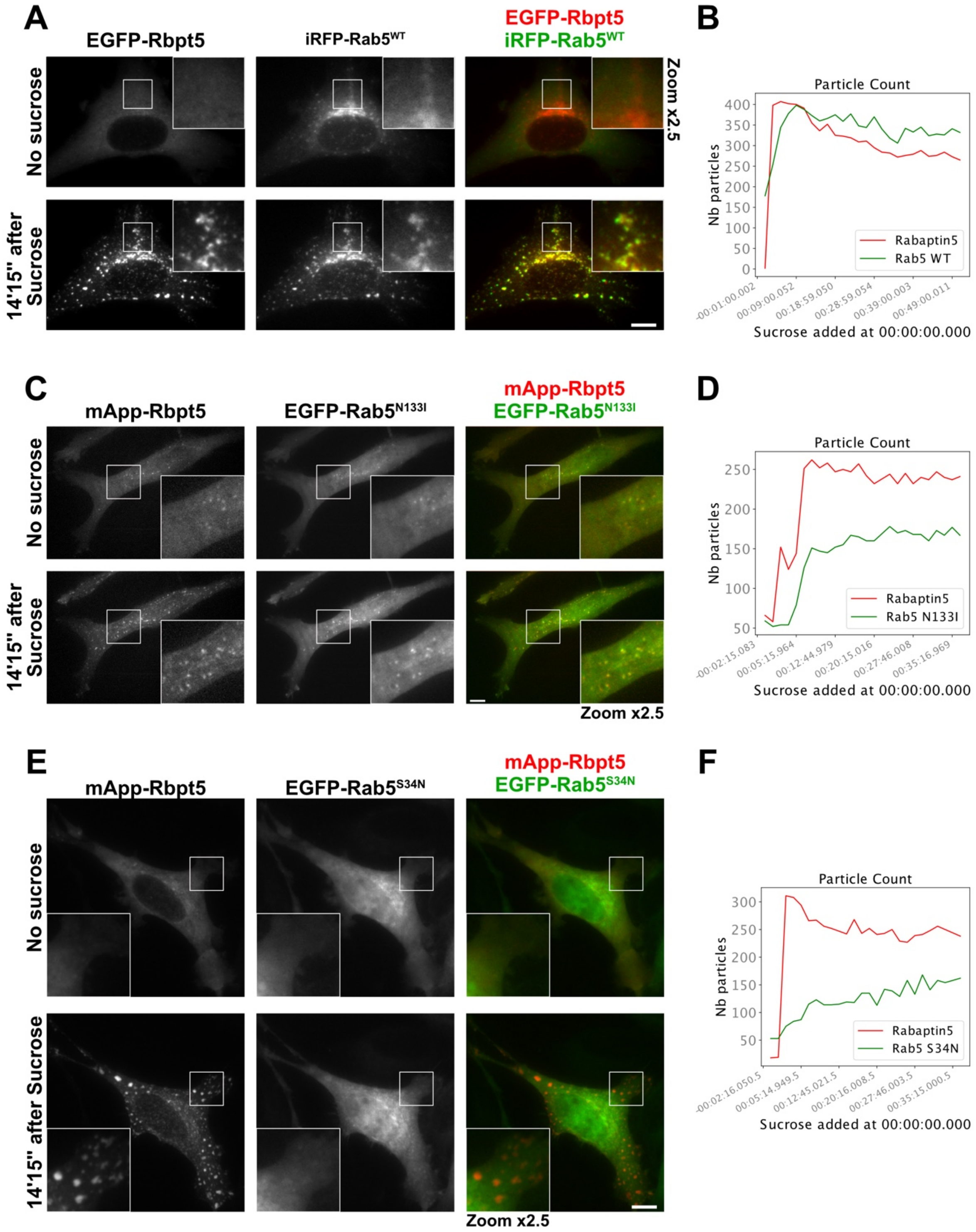


**Figure S2. Rab5 mutations impairing prenylation or GTP binding also prevent Rab5 from enriching in Rabaptin5 condensates.**

Time-lapse imaging of cells co-expressing Rabaptin5 and Rab5 and respective particle quantification (**A,B-** WT, **C,D-** prenylation deficient N133I, or **E,F-** GTP binding deficient S34N) reveals decreased Rab5 enrichment on Rabaptin5 condensates as well as a decrease in the number of observed puncta in Rab5 mutants. Scale bars: 10µm.

**Figure S3**

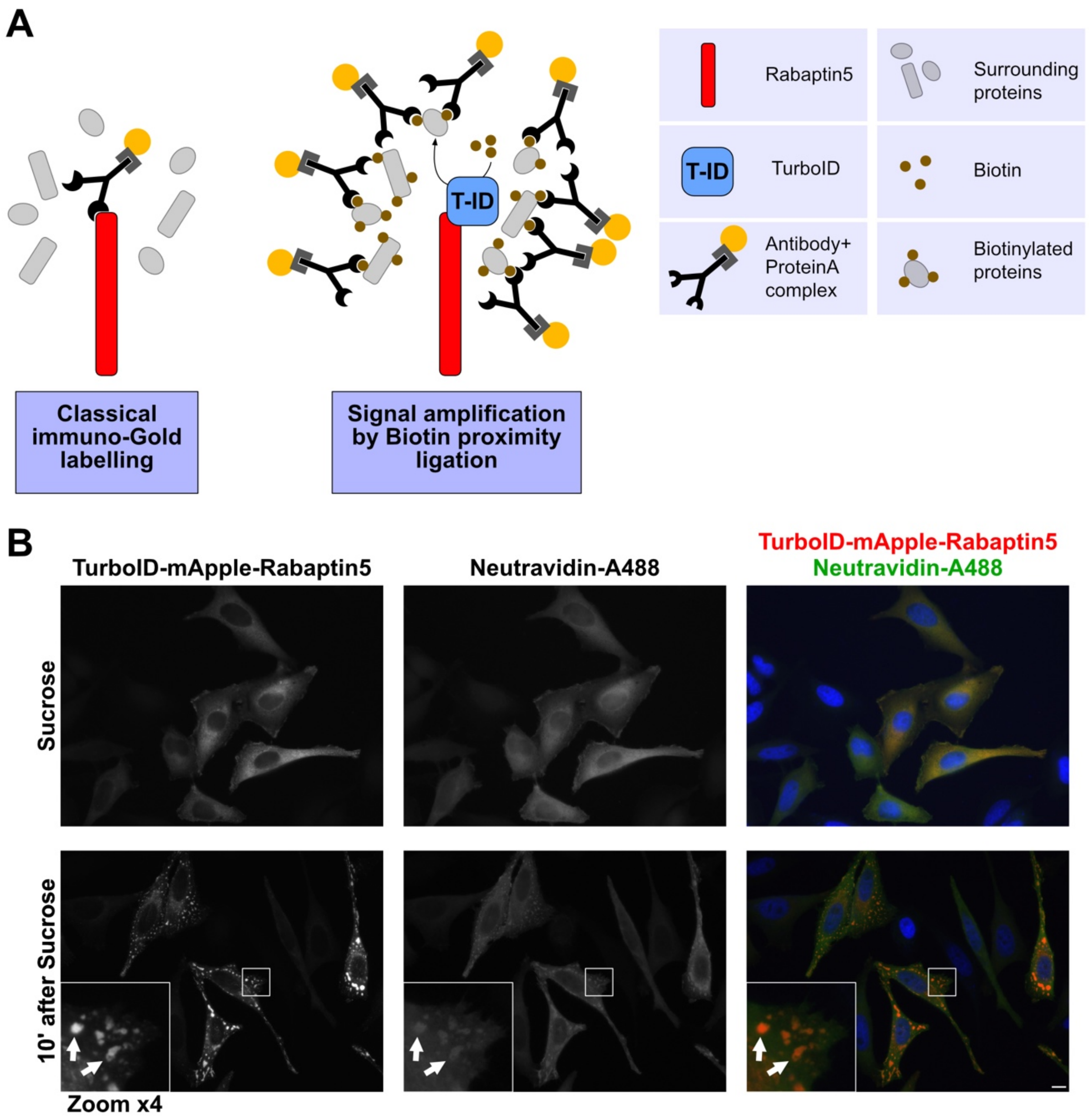


**Figure S3. Upon treatment with sucrose and biotin, TurboID-mApple-Rabaptin5 is able to efficiently biotinylate its surroundings within condensates.**

**A-** Scheme illustrating the use of Rabaptin5 fused to the promiscuous biotin ligase Turbo-ID, which allows biotinylation of proteins in Rabaptin5 surroundings and allows ImmunoGold signal amplification. **B-** HeLa cells were transfected with TurboID-mApple-Rabaptin 5 and treated with sucrose (147µM) in presence of biotin (40µM) for 10 min, then washed and fixed. Cells were then processed for cryo-immuno-electron microscopy (Fig. 3D) or stained using neutravidin-A488. White arrows point at biotinylated condensates. Scale bar : 10µm.

Figure S4

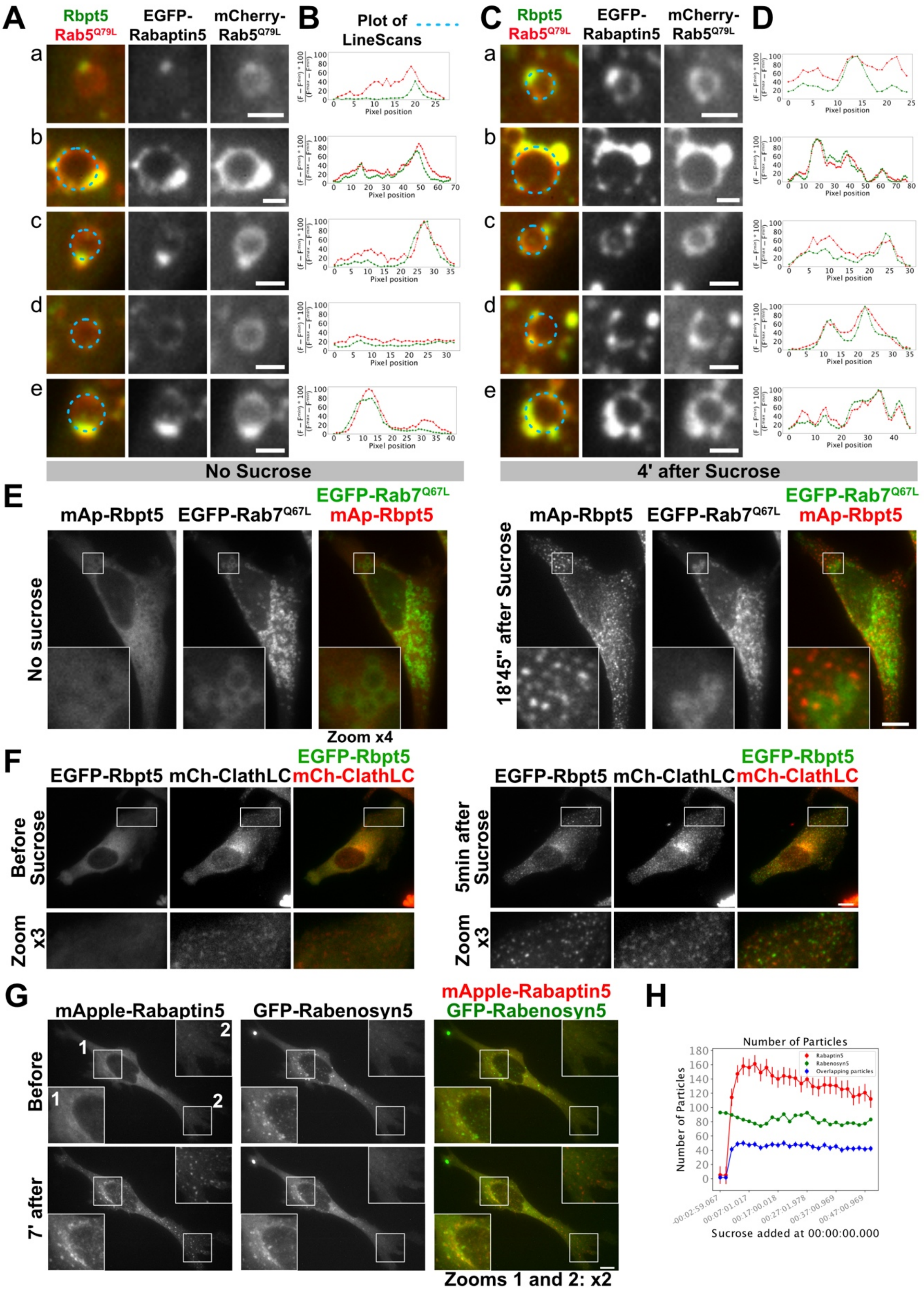


**Figure S4. Rabaptin5 condensates localize close to membranes and is accompanied by an enrichment in Rab5 but do not co-localize with markers of mature early endosomes nor with early markers of endocytosis**

**A-D** Cells transfected with EGFP-Rabaptin5 and mCherry-Rab5 Q79L mutant present Rabaptin5 condensates docked on enlarged endosomes, capable of growth and Rab5 enrichment upon sucrose treatment. **A,C-** Close-up on several enlarged endosomes followed before and after sucrose. Dashed blue curves marking the perimeter scans plotted in **B** and **D. B,D-** Straighten perimeter scans of enlarged endosomes before and after sucrose respectively. Like in Figure 3, intensity profiles were normalized using the minimum between the minimum value found in profile before and after sucrose, and similarly for the maximum, for each channel independently. **E-** Cells transfected mApple-Rabaptin5 and EGFP-Rab7 Q67L mutant does not present docking of Rabaptin5 condensates on Rab7 positive enlarged endosomes. **F-** Co-expression of Rabaptin5 and clathrin light-chain reveals Rabaptin5 condensate appearance and some increase in intensity on clathrin puncta, as well as some juxtaposition between the two markers but no co-localization. **G-** Time-lapse imaging of cells co-expressing mApple-Rabaptin5 and GFP-Rabenosyn5 condensates reveals juxtaposition but not co-localization between the two proteins upon sucrose addition. Zoom 1 shows central particles co-labeled with Rabaptin5 and Rabenosyn5. Zoom 2 shows newly showing peripheral particles labeled with Rabaptin5 and little or no Rabenosyn5. **H-** Puncta quantification shows particle number increase for Rabaptin5 but not for Rabenosyn5. It also reveals little overlapping between Rabenosyn5 and Rabaptin5 puncta. Scale bars E-G: 10µm; scale bars A,C: 2µm.

**Figure S5**

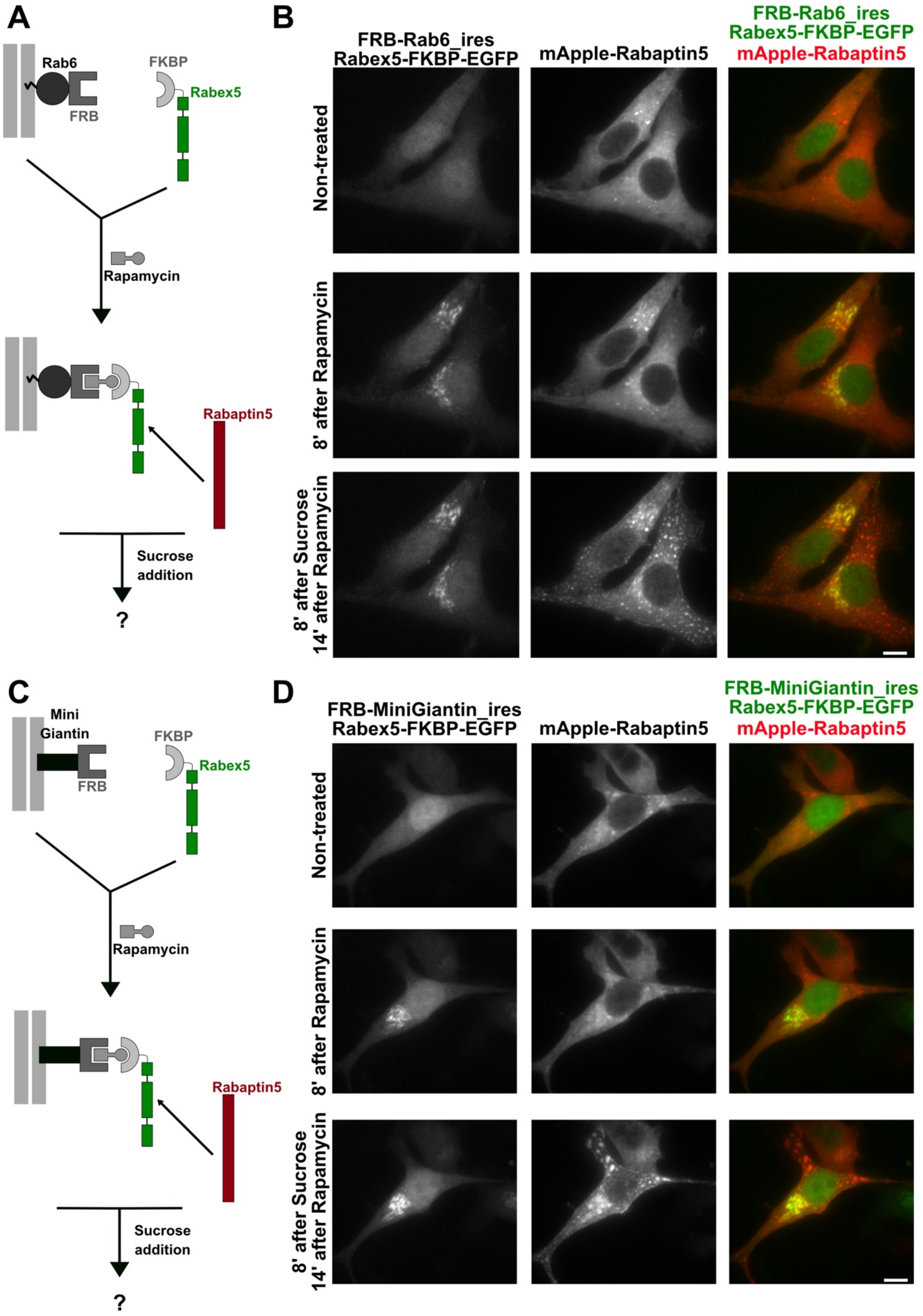


**Figure S5. Artificial indirect recruitment of Rabaptin5 on "foreign" membranes potentiate Rabaptin5 condensate formation.**

**A,C-** Experiment layouts. **B,D-** Rabaptin5 artificially recruited at the Golgi is able to nucleate condensates . Scale bars: 10µm.

**Figure S6**

**A**

| | Length | % identity with H. sapiens Isoform1 | % Similarity with H. sapiens Isoform1 | Gaps compared to H. sapiens Isoform1 |
|---|---|---|---|---|
| Homo sapiens Isoform1 | 862 | | | |
| Rattus norvegicus | 862 | 95,94% | 98,03% | 0% |
| Mus musculus isoform1 | 862 | 96,29% | 98,26% | 0% |
| Bos taurus | 862 | 97,22% | 98,96% | 0% |
| Gallus gallus | 860 | 88,05% | 93,62% | 0,23% |
| Xenopus tropicalis | 853 | 77,73% | 88,40% | 1,04% |
| Danio renio isoform1 | 850 | 71,07% | 83,47% | 2,13% |
| Drosophila melanogaster | 634 | 24,92% | 45,43% | 14,04% |

**B**

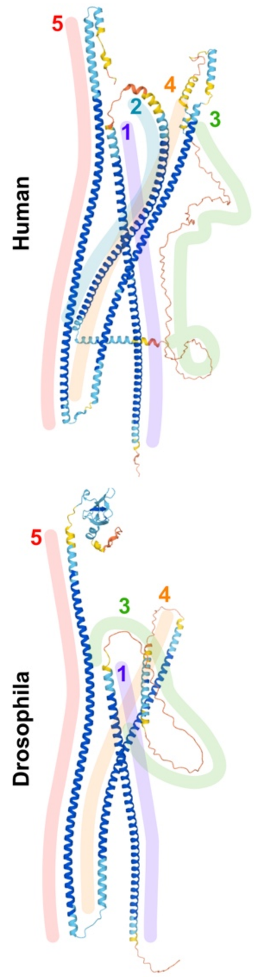


**C**

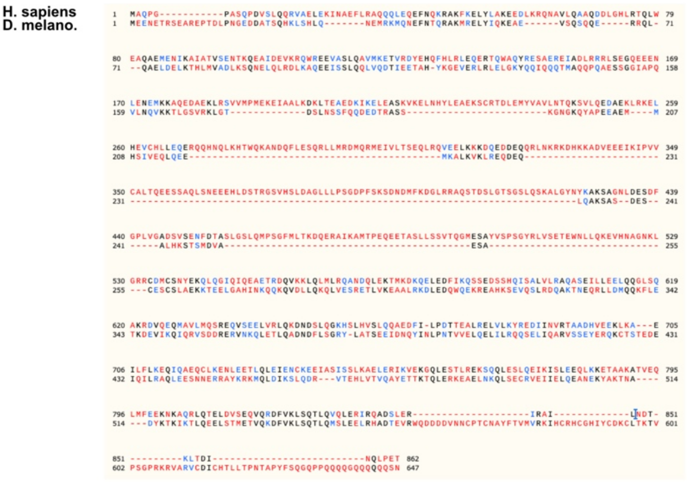


**D**

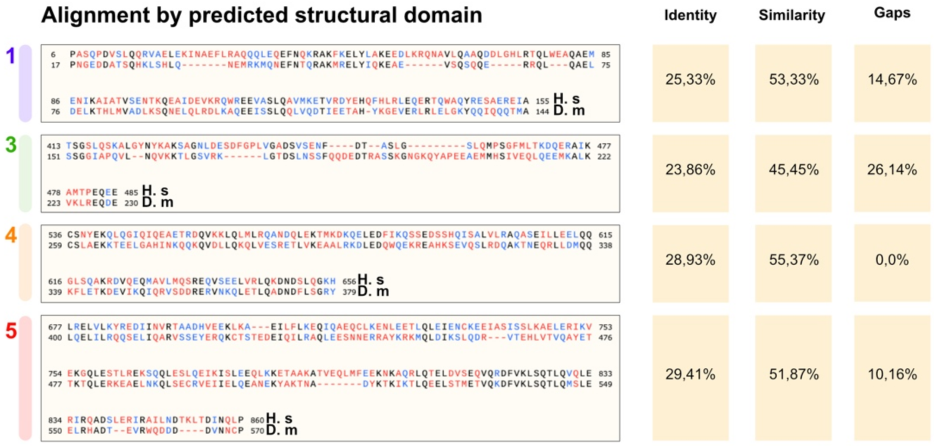


**Figure S6. Rabaptin5 is evolutionary conserved in sequence among animals and in structure compared to insects.**

**A-** Table shows the amino acid sequence identities or similarities between Human Rabaptin5 and other species versions. **B-** Alphafold 3D structure prediction of Human and Drosophila versions. **C-** Overall alignment of amino-acid sequences (SnapGene). **D-** Alignment region by region defined by structure prediction in **B**.

**Figure S7**

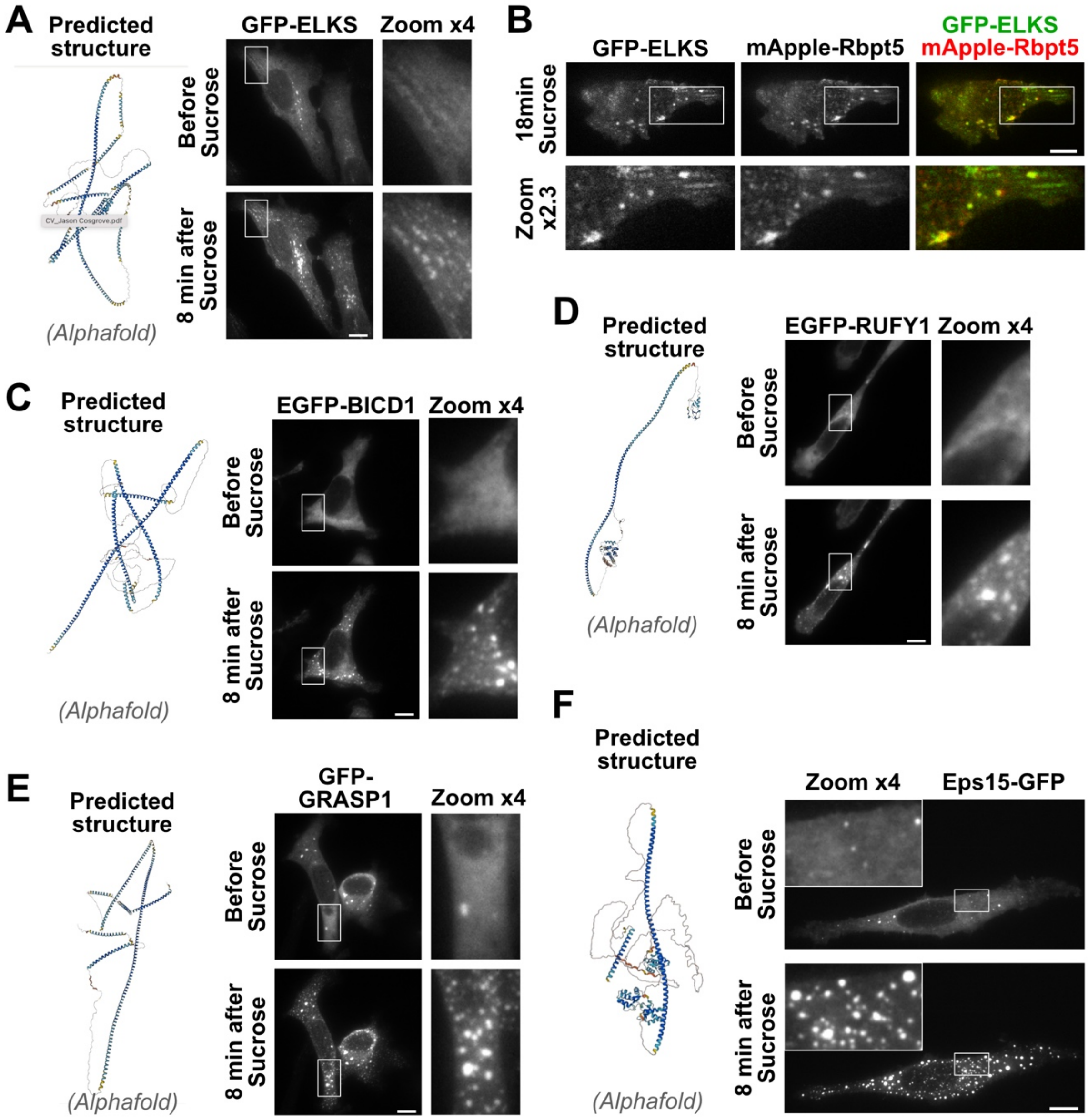


**Figure S7. Other Rab partners presenting common 3D features with Rabaptin (long alpha-helixes and long unstructured domains) are also able to undergo LLPS.**

Cells were transfected with indicated Rab partners and treated with medium supplemented with 147mM sucrose. **A,C-F-** Alphafold predicted structures and pictures before and after sucrose are shown, for ELKS, BICD1, RUFY1 (a.k.a. Rabip4'), GRASP1 and Eps15 respectively. **B-** Cells co-expressing mApple-Rabaptin5 and EGFP-ELKS present partial co-localisation. Scale bars: 10µm

### Movie Legends

**Movie S1. Rabaptin5 condensates forming upon Sucrose addition are reversible.**
HeLa cells were transfected with EGFP-Rabaptin5 24h before the acquisition with a 2min time-lapse under a spinning disk microscope. Sucrose was added at 2, 22 and 42 minutes, and respective wash-outs were performed at 13, 33 and 53 minutes.

**Movie S2. Rabaptin5 condensates are able to undergo fusion.**
HeLa cells were transfected with EGFP-Rabaptin5 24h before the acquisition with a 1 second time-lapse under a spinning disk microscope. Sucrose was added at 18 seconds after the start of the movie.

**Movie S3. Rabaptin5 condensates can appear at the edges of the cell.**
HeLa cells were transfected with EGFP-Rabaptin5 24h before the acquisition with a 1 second time-lapse under a spinning disk microscope. Sucrose was added at 20 seconds after the start of the movie.

**Movie S4. Fusion of nested Rabaptin5 and Eps15 condensates is homotypic.**
HeLa cells were transfected with mApple-Rabaptin5 and Eps15-GFP 24h before the acquisition with a 1 second time-lapse under a spinning disk microscope. Sucrose was added at 28 seconds after the start of the movie. Several events of fusion between nestled condensates are observable.
